# The Rab11-Family Interacting Proteins reveal selective interaction of mammalian recycling endosomes with the *Toxoplasma* parasitophorous vacuole in a Rab11- and Arf6-dependent manner

**DOI:** 10.1101/2021.06.01.446625

**Authors:** Eric J. Hartman, Julia D. Romano, Isabelle Coppens

## Abstract

After invasion of mammalian cells, the parasite *Toxoplasma gondii* multiplies in a self-made membrane-bound compartment, the parasitophorous vacuole (PV). We previously showed that intravacuolar *Toxoplasma* interacts with many host cell organelles, especially recycling endosomes, and further manipulates the host endocytic recycling through the sequestration of Rab11 vesicles into the PV. Mammalian Rab-PV interactions are likely mediated by *Toxoplasma* and host proteins that remain to be identified. In this context, we have examined the specificity of host Rab vesicle interaction with the PV by monitoring the recruitment of subtypes of Rab11 vesicles differing in their composition in Rab11-Family Interacting Proteins (FIPs). We found that vesicles with FIPs from Class I (FIP1C, FIP2, FIP5) or Class II (FIP3, FIP4) are distributed at the PV and detected to varying degrees inside the PV. The PV delivery of vesicles with FIPs from Class I, but not Class II, is Rab11-dependent. In addition to Rab11, FIP3 binds to Arf6, and vesicles associated with FIP3-Arf6 complexes are observed within the PV. Binding of FIP3 to either Rab11 or Arf6 significantly increases the internalization of vesicles into the PV. These data point to a selective process of host recycling endosome recognition and scavenging mediated by *Toxoplasma*.

## Introduction

The small monomeric Rab GTPases belong to the Ras superfamily and are master regulators of vesicular trafficking and signaling pathways (Chavrier and Goud, 1999). Many intracellular pathogens hijack the host Rab membrane trafficking machinery to thrive in a hostile environment. Among protozoan parasites, Apicomplexa are obligate intracellular microbes that invade mammalian cells by creating a protective, nonfusogenic membrane-bound compartment, the parasitophorous vacuole (PV). Recent evidence demonstrates that apicomplexan parasites manipulate several mammalian Rab vesicle trafficking pathways to scavenge nutrients or to exploit the functions of Rab GTPase proteins for immune evasion (reviewed in Coppens and Romano, 2020). The apicomplexan *Toxoplasma gondii*, the etiologic agent of toxoplasmosis, diverts many host Rab vesicles to the PV and sequesters some of them in the PV lumen through deep invaginations of the PV membrane (Romano *et al*., 2013; 2017; Nolan *et al*., 2017; Coppens and Romano, 2020). *Toxoplasma* predominantly hijacks the GTP-bound form of host Rab proteins to intercept vesicular traffic as a means for lipid procuration. Indeed, the parasite salvages sphingolipids manufactured in the host ER/Golgi and accesses these lipids through the interception of Rab14, Rab30 and Rab43 vesicles (Romano *et al*., 2013).

The Rab11 proteins, comprising Rab11A, Rab11B and Rab11C/Rab25, localize to the *trans*-Golgi network (TGN), post-Golgi vesicles and pericentriolar recycling endosomes (RE), and they function in the recycling of material to the plasma membrane via RE (Welz *et al*., 2014; Grant and Donaldson, 2009). Rab11 proteins regulate the recycling of many receptors and adhesion proteins to the cell surface and play roles in diverse cellular functions, including ciliogenesis, cytokinesis, neuritogenesis and oogenesis (Kelly *et al*., 2012). Abnormal Rab11 activity (deficiency or overexpression) has been shown to be implicated in the progression of numerous human diseases (e.g., cancers, neurodegeneration, diabetes) (Bhuin and Roy, 2015). Rab11A, Rab11B and Rab11C/Rab25 localize to different cellular compartments, suggesting division of selective functions between the three subfamily members. Rab11A is involved in the targeted delivery of material to the cleavage furrow/midbody during cell division, phagocytosis, long-term potentiation of synaptic transmission, and cell migration (Horgan *et al*., 2012; Cox *et al*., 2000; Wang *et al*., 2008; Kelly *et al*., 2011; Yoon *et al*., 2005; Caswell *et al*., 2008) while Rab11B is essential for the recycling of the transferrin receptor (TfR) (Schlierf *et al*., 2000), the epithelial sodium channel, the cystic fibrosis transmembrane conductance regulator, for the regulation of the trafficking of vacuolar type H^+^-ATPases, and for calcium-induced exocytosis (Khvotchev *et al*., 2003; Silvis *et al*., 2009; Sugawara *et al*., 2009; Best *et al*., 2011; Butterworth *et al*., 2012). Rab11C/Rab25 is associated with apical RE and regulates the processes of transcytosis and plasma membrane recycling (Casanova *et al*., 1999). Among mammalian Rab vesicles, those from the recycling and secretory (anterograde) pathways are detected in the vast majority of the *Toxoplasma* PV (Romano *et al*., 2017). In particular, 89% and 100% of PV contain Rab11A and Rab11B vesicles, respectively.

Rab11 proteins interact with different effectors depending on their GTP/GDP status that is regulated by guanine nucleotide exchange factors (GEF) and GTPase-activating proteins (GAP) (Zerial and McBride, 2001). In the active state, Rab11 proteins bind to effector proteins termed the Rab11-Family Interacting Proteins (FIPs), allowing Rab11 proteins to recruit cellular motor proteins (Horgan and McCaffrey, 2009). FIPs interact indiscriminately with GTP-bound Rab11A and Rab11B via a conserved 20-aa C-terminal domain, named the Rab11-binding domain (RBD) (Junutula *et al*., 2004; Horgan and McCaffrey, 2009; Eathiraj *et al*., 2006; Jagoe *et al*., 2006). Mutations introduced into the RBD of FIP disrupts binding to Rab11, and therefore alters the localization of the FIP (Lindsay and McCaffrey 2004a; Junutula *et al*., 2004; Jagoe *et al*., 2006; Wilson *et al*., 2005; Fielding *et al*., 2005; Horgan *et al*., 2007; Meyers and Prekeris, 2002). In addition to the RBD, each FIP has an α-helical coiled-coil structure mediating FIP homodimerization necessary to execute FIP cellular functions (Wei *et al*., 2006; 2009; Horgan *et al*., 2007). FIPs form heterotetrameric complexes with Rab11 composed of two Rab11 and two FIP molecules. Based on their primary structure, FIPs are subcategorized into class I, with FIP1C (*alias* Rab-coupling protein), FIP2 and FIP5 (*alias* Rip11, Gaf-1/Gaf-1b, pp75 or gamma-SNAP Associated Factor), and class II, with FIP3 (*alias* Arfophilin or Eferin) and FIP4 (*alias* Arfophilin-2). Class I FIPs have a phospholipid-binding C2-domain and are involved in many endosomal recycling and trafficking processes (Lindsy *et al*., 2002; Lindsay and McCaffrey 2004b; Fan *et al*., 2004; Prekeris *et al*., 2000). Class II FIPs contain a calcium binding EF-Hand domain, function in targeting new membranes to the cleavage furrow for the completion of cytokinesis and are necessary for the structural integrity of the pericentrosomal endocytic recycling compartment (ERC) (Horgan *et al*., 2007). The ERC is constituted of a large complex of heterogeneous subsets of RE, tubular RE and small transport intermediates (Maxfield and McGraw, 2004). FIP3 and FIP4 also bind members of the Arf GTPase family, most notably Arf6, on an Arf-binding domain (ABD) in the C-terminal region that is distinct from the RBD (Shin *et al*., 1999; Fielding *et al*., 2005; Shiba *et al*., 2006). Arf6 forms ternary complexes with Rab11 and FIP3 or FIP4, and mediates the recruitment of Rab11-FIP3 and Rab11-FIP4 to the cleavage furrow (Horgan *et al*., 2004; Wallace *et al*., 2002; Wilson *et al*., 2004; Fielding *et al*., 2005). Deletion of the ABD from FIP3 in neurons leads to defects in Arf6-regulated delivery of endosome recycling pathways, such as aberrant cytoplasmic retention of N-cadherin and syntaxin 12. The resulting phenotype is impairment in neuron migration, which reveals the importance of Arf6-FIP endosomal trafficking pathways (Hara *et al*., 2016).

In this study, we sought to address the specificity of host Rab vesicle interaction with the PV of *Toxoplasma* by focusing on mammalian Rab11 vesicles. In particular, we sought to examine whether *Toxoplasma* hijacks indiscriminately any host Rab11A or Rab11B vesicle regardless of composition or localization in infected cells, or if the parasite selectively recognizes subpopulations of Rab11A or Rab11B vesicles associated with specific effectors. Using quantitative microscopy, we assessed the distribution of ectopically expressed, fluorescently tagged mammalian FIP (FIP1C, FIP2, FIP3, FIP4 and FIP5) and Arf6 in *Toxoplasma*-infected cells. We further determined the contribution of Rab11 or Arf6 binding to the internalization of FIP3 into the PV through mutations introduced in the RBD and/or ABD of FIP3. Our *in vitro* assays reveal differential interception and PV internalization of host RE based on effector and modulator protein composition.

## Results

### Mammalian Rab11 vesicle subpopulations with either Class I or Class II FIPs are associated with the PV of *Toxoplasma*

In mammalian cells, the intravacuolar parasite *Toxoplasma* intercepts the trafficking of host recycling vesicles (Romano *et al*., 2017). The recruitment of host Rab11 vesicles at the PV was monitored by immunofluorescence assays (IFA) in infected VERO cells using an antibody recognizing Rab11 proteins (Figure 1A). Compared to uninfected cells showing Rab11 puncta near the nucleus and dispersed throughout the cytoplasm, the Rab11 signal in infected cells was observed surrounding the PV (delineated by TgGRA7 staining). The interior of the PV is characterized by the presence of an entangled network of membranous tubules (∼30 nm diameter), named the IntraVacuolar Network (IVN) (Sibley *et al*., 1995). Previously, we illustrated by electron microscopy (EM) the attachment of IVN tubules to the PV membrane providing ‘open gates’ for the entry of host material into the PV, and we detected host organelles, including Rab vesicles within the tubules forming the IVN (Romano *et al*., 2017). Our IFA on infected cells confirmed the intra-PV distribution of Rab11 vesicles at the IVN stained with TgGRA7 (Figure 1A). In VERO cells expressing GFP-Rab11A or GFP-Rab11B, the fluorescent signal was perinuclear in uninfected cells then redistributed at the PV upon infection, with GFP foci also observed in the PV lumen (Figure 1B and 1C). Confirmation of the GFP signal corresponding to Rab11A in transfected cells was further confirmed by immunostaining with anti-Rab11A antibody, showing significant colocalization of the two fluorescent signals in uninfected VERO cells (Figure 1D, panel a). In transfected and infected VERO cells, the GFP and antibody signals were observed around the PV (Figure 1D, panel b) and within the vacuole (Figure 1D, panel c).

**Figure 1.**
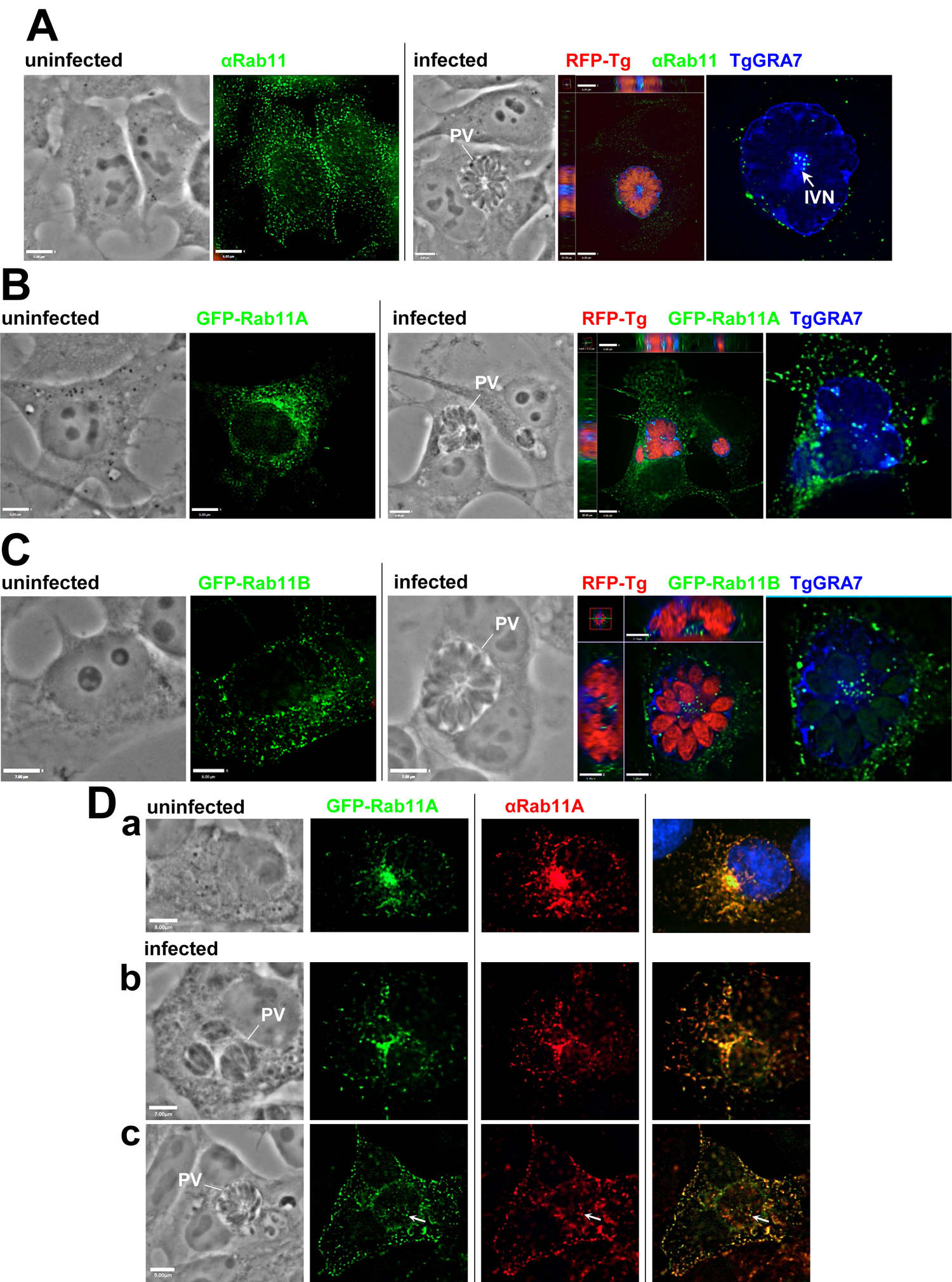
Distribution of mammalian Rab11 vesicles in *Toxoplasma*-infected cells. **A.** Fluorescence microscopy of VERO cells, uninfected or infected with RFP-expressing *Toxoplasma* (RFP-Tg) for 24h, before immunostaining with anti-Rab11 and anti-TgGRA7 (PVM/IVN) antibodies. For all images, individual z-slices and orthogonal views highlighting the PV, are shown. **B-C.** Fluorescence microscopy of VERO cells expressing GFP-Rab11A (in B) or GFP-Rab11B-(in C), uninfected or infected with RFP-expressing *Toxoplasma* for 24h, before immunostaining with anti-TgGRA7 (PVM/IVN) antibody. **D.** Fluorescence microscopy of VERO cells expressing GFP-Rab11A-, uninfected (panel a) or infected with *Toxoplasma* for 24h, before immunostaining with anti-Rab11A antibody, showing colocalization of the Rab11A signals surrounding the PV (panel b) and inside the PV (panel c).

Next, we wanted to examine whether *Toxoplasma* selectively recruits subpopulations of mammalian Rab11 vesicles associated with specific effectors that govern their localization and recycling function. We transiently transfected HeLa cells with a plasmid containing a FIP from Class I (FIP1C, FIP2, FIP5) or Class II (FIP3, FIP4) fused N-terminally with GFP. Post-transfection, cells were infected with RFP-expressing *Toxoplasma* for 24h. In uninfected HeLa cells, GFP-FIP1C exhibited a wide distribution in the cytoplasm demarcating tubular structures preferentially distributed at the perinuclear region (Figure 2A, panel a). In infected cells, GFP-FIP1C structures were observed concentrated around the PV. In uninfected cells expressing GFP-FIP2, the fluorescent pattern was similar to that of GFP-FIP1C, and upon infection, it also delocalized to the PV (Figure 2A, panel b). Compared to the GFP-FIP1C and GFP-FIP2, the fluorescent signal of GFP-FIP5 was predominantly centralized in the juxtanuclear region of uninfected cells (Figure 2A, panel c). Following infection, GFP-FIP5 localization was more prominent at the PV than at the host nucleus.

**Figure 2.**
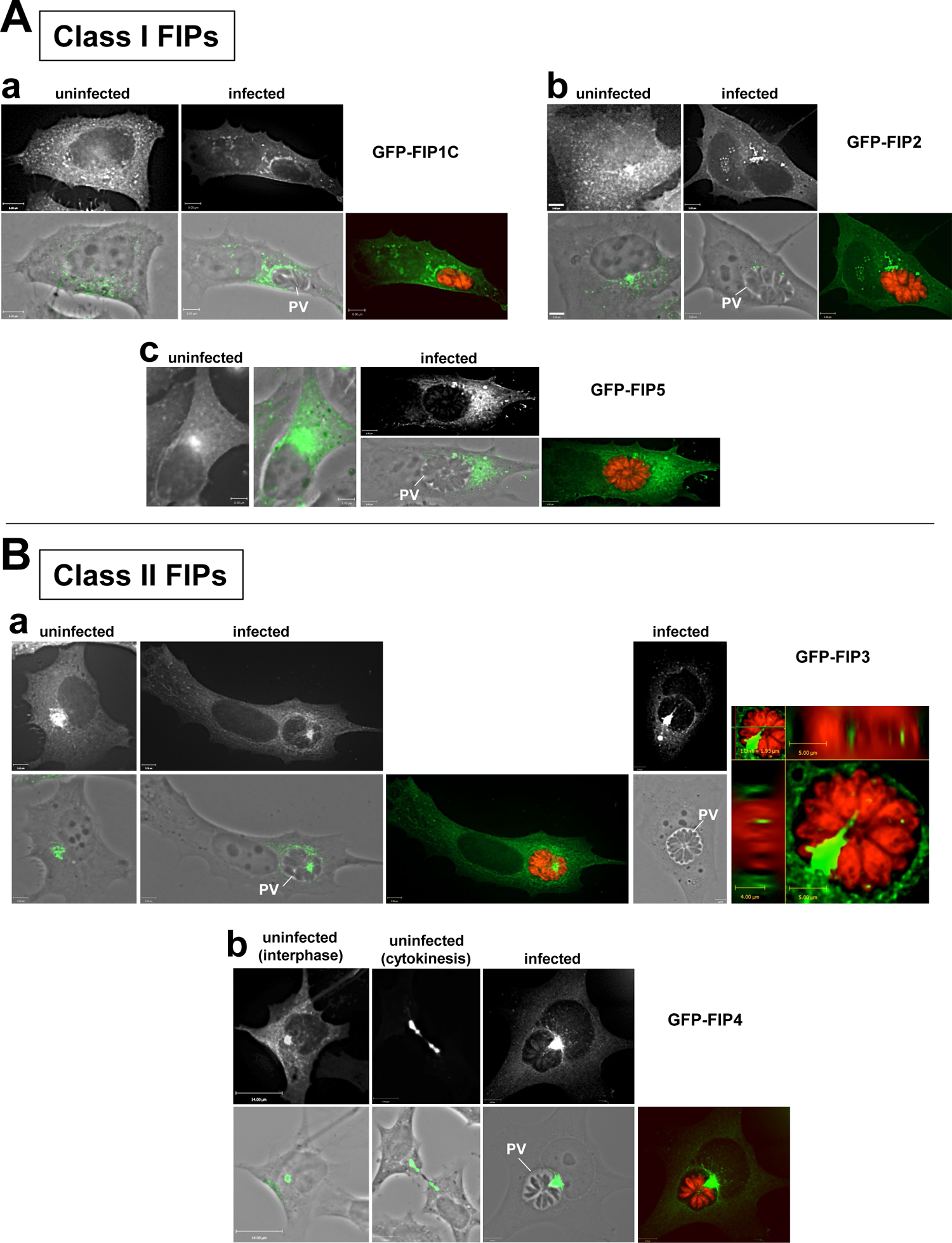
Distribution of mammalian vesicles containing FIP from class I and II in *Toxoplasma*-infected cells. **A-B.** Fluorescence microscopy of HeLa cells transfected with Class I FIPs (GFP-FIP1C, GFP-FIP2 or GFP-FIP5 in A) and Class II FIPs (GFP-FIP3 or GFP-FIP4 in B), uninfected or infected with RFP-expressing *Toxoplasma* for 24h. Phase contrast overlaid with the GFP-FIP signal is shown as well as the GFP-FIP alone and with the RFP signal.

Next, we compared the localization of Class II FIPs (FIP3, FIP4) in uninfected and infected cells. It has been reported that exogenous expression of FIP3 or FIP4, which localize to the endocytic recycling compartment (ERC), induces alteration in the morphology of the ERC as this organelle condenses around the microtubule-organizing center (MTOC) at the pericentrosomal region of cells during interphase (Horgan *et al*., 2004; Wallace *et al*., 2002). During cytokinesis, FIP3 and FIP4 vesicles move to the cleavage furrow, a process also amplified upon FIP3 and FIP4 overexpression (Horgan *et al*., 2004; Wallace *et al*., 2002; Wilson *et al*., 2005; Fielding *et al*., 2005). In HeLa cells expressing GFP-FIP3 or GFP-FIP4, we confirmed an intense fluorescent staining at the ERC and the cleavage furrow in non-dividing and mitotic cells, respectively (Figure 2B, panels a and b). Upon *Toxoplasma* infection, the GFP-FIP3 or GFP-FIP4-positive ERC was delocalized from its pericentrosomal location and associated with the PV.

### *Toxoplasma* internalizes mammalian Rab11 vesicle subpopulations with either Class I or Class II FIPs into the PV with some selectivity

Aggregation at the *Toxoplasma* PV of host vesicles marked with FIP1C, FIP2, FIP5, FIP3 or FIP4 may precede the internalization of these vesicles into the PV. In infected HeLa cells expressing each GFP-FIP protein, the intra-PV localization of GFP-FIP foci was monitored by fluorescence microscopy by collecting a series of optical z-sections throughout the PV defined by TgGRA7 immunostaining. GFP-FIP foci were observed within the PV for all the FIP constructs from Class I (Figure 3A) and Class II (Figure 3B). However, the percentage of PV positive for intra-PV GFP-FIP foci was different among the FIPs; the lowest PV percentage was measured for FIP1C (12%) and the highest for FIP3 (88%) (Figure 3C). Approximately 50% of PV contained FIP2, FIP5 or FIP4 vesicles. A *Toxoplasma* mutant lacking two-IVN-localized proteins involved in IVN biogenesis (TgGRA2 and TgGRA6) shows complete disruption of this network, which is reduced to loose membrane whorls (Mercier *et al*., 2002) and is impaired in host organelle sequestration into the PV (Figure S1; Romano *et al*., 2017). This mutant was used as a negative control in our assays, and as expected, no GFP foci were detected in the vast majority of the Δgra2Δgra6 PV in HeLa cells expressing GFP-FIPs from Class I and Class II (Figure S2).

**Figure 3.**
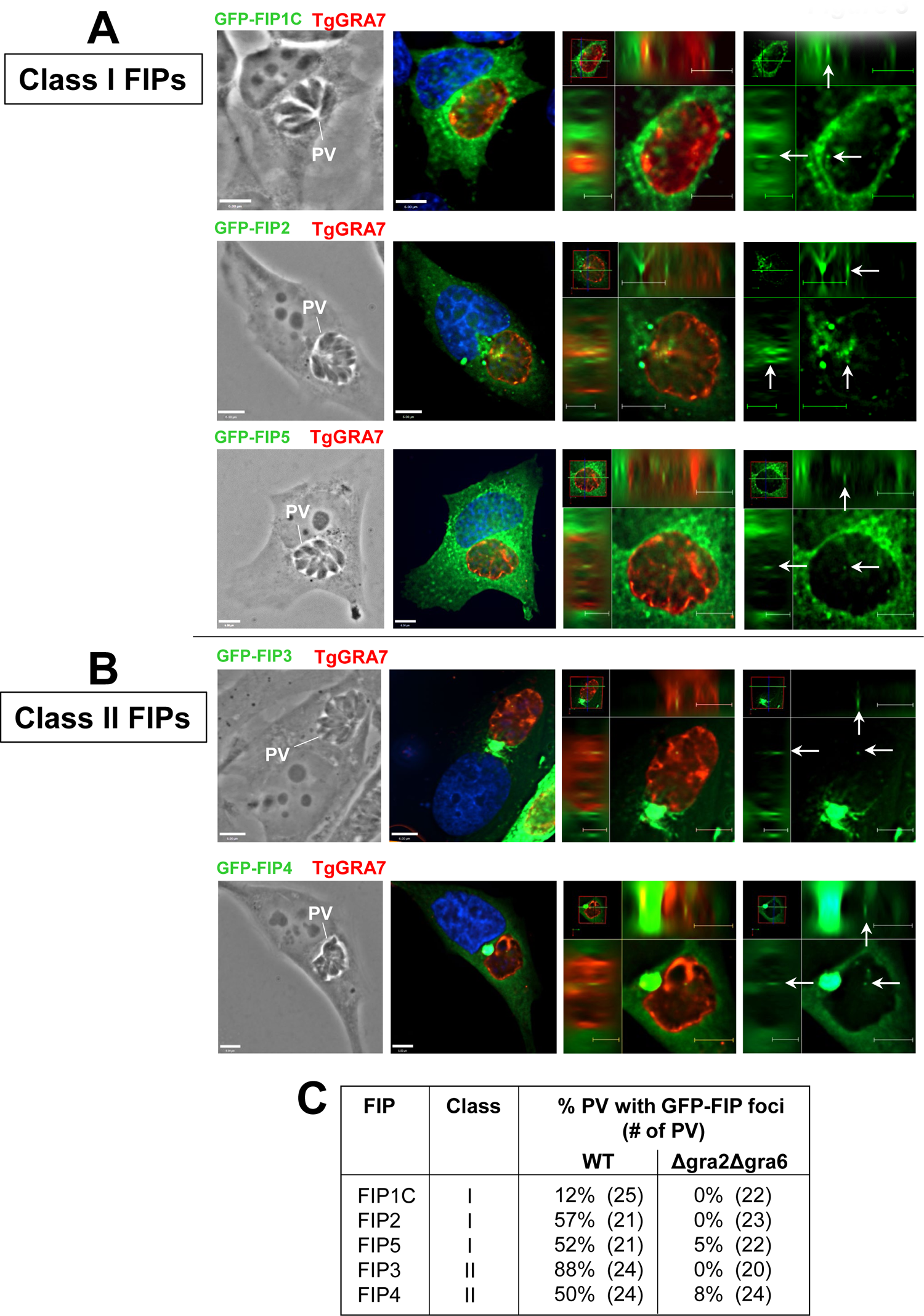
Distribution of mammalian vesicles containing FIP from class I and II within the *Toxoplasma* PV. **A-B.** Fluorescence microscopy of HeLa cells transfected with GFP-FIP (Class I in A: GFP-FIP1C, GFP-FIP2 or GFP-FIP5; Class II in B: GFP-FIP3 or GFP-FIP4), uninfected or infected with RFP-expressing *Toxoplasma* for 24h before immunostaining with anti-TgGRA7 antibody. DAPI in blue. For all images, individual z-slices and orthogonal views are shown. Arrows show intra-PV GFP-FIP foci. **C.** Quantification of the internalization of Class I and II FIP vesicles within the *Toxoplasma* PV. Percent of PV with intra-PV foci was assessed based on PV viewed in A and B. Negative controls include PV from Δgra2Δgra6 mutants impaired in host vesicle internalization.

### PV sequestration of vesicles marked with Class I FIP but not Class II FIP requires the formation of Rab11-FIP complexes

FIP proteins play a role in targeting Rab11 to different endocytic compartments by competing with each other for Rab11 binding (Meyers and Prekeris, 2002). We next wanted to examine whether the recruitment of FIP-associated vesicles at the PV and their internalization into the vacuole require the presence of Rab11 bound to FIP. Substitution of the hydrophobic isoleucine residue to a negatively charged glutamic acid within the Rab11-binding domain (RBD) of FIP1C (I621E mutation), FIP2 (I481E mutation), FIP5(I630E mutation), and FIP3 (I738E mutation) or replacement of the tripeptidic sequence tyrosine-methionine-aspartate by three alanine residues in the RBD of FIP4 (YMD617-619AAA) totally abrogates FIP binding to Rab11 and the distribution of FIP-RBD mutant vesicles is grossly altered in mammalian cells (Lindsay and McCaffrey 2004a; Junutula *et al*., 2004; Jagoe *et al*., 2006; Wilson *et al*., 2005; Fielding *et al*., 2005; Horgan *et al*., 2007; Meyers and Prekeris, 2002). We transiently transfected HeLa cells with plasmids containing each FIP-RBD mutant fused to GFP, and the GFP signal was predominantly dispersed throughout the cell, with a noticeable loss of pericentrosomal ERC staining for FIP3 and FIP4 (Figure 4A and 4B). Following a 24h infection of transfected cells, fluorescent perivacuolar labeling was observed, suggesting the ability of the parasite to recruit FIP-RDB mutant vesicles, even those scattered in the host cytoplasm. Interestingly, many PV in HeLa cells expressing the mutants FIP3-RBD and FIP4-RBD contained numerous GFP foci (Figure 4B). In contrast, no intra-PV GFP signal was detected for the Class I FIP-RBD mutants (Figure 5A). For the Class II mutants, all PV were positive for FIP4-RBD mutant vesicles and three-fourths of PV for FIP3-RBD mutant vesicles (Figure 5B). No GFP foci of FIP3-RBD and FIP4-RBD mutants were spotted in the PV of Δgra2Δgra6 parasites, as expected (Figure S2). The number of intra-PV GFP-FIP foci was counted for all PV in infected HeLa cells expressing Class I and II FIP-WT and Class II FIP-RBD mutants (Figure 6A). Examining the statistical significance for all PV, including ones without intra-PV foci better represents the accuracy of the percent of positive PV counts, thus revealing how often internalization events occur. Focusing on the number of foci only present in positive PV highlights any differences in intra-PV foci abundance; our data in Figure 6B show that the greatest number of intra-PV vesicles was scored for the Class II FIP mutants, with an average number of 7 foci (7.4 FIP3-RBD foci/positive PV) and (6.3 FIP4-RBD foci/positive PV).

**Figure 4.**
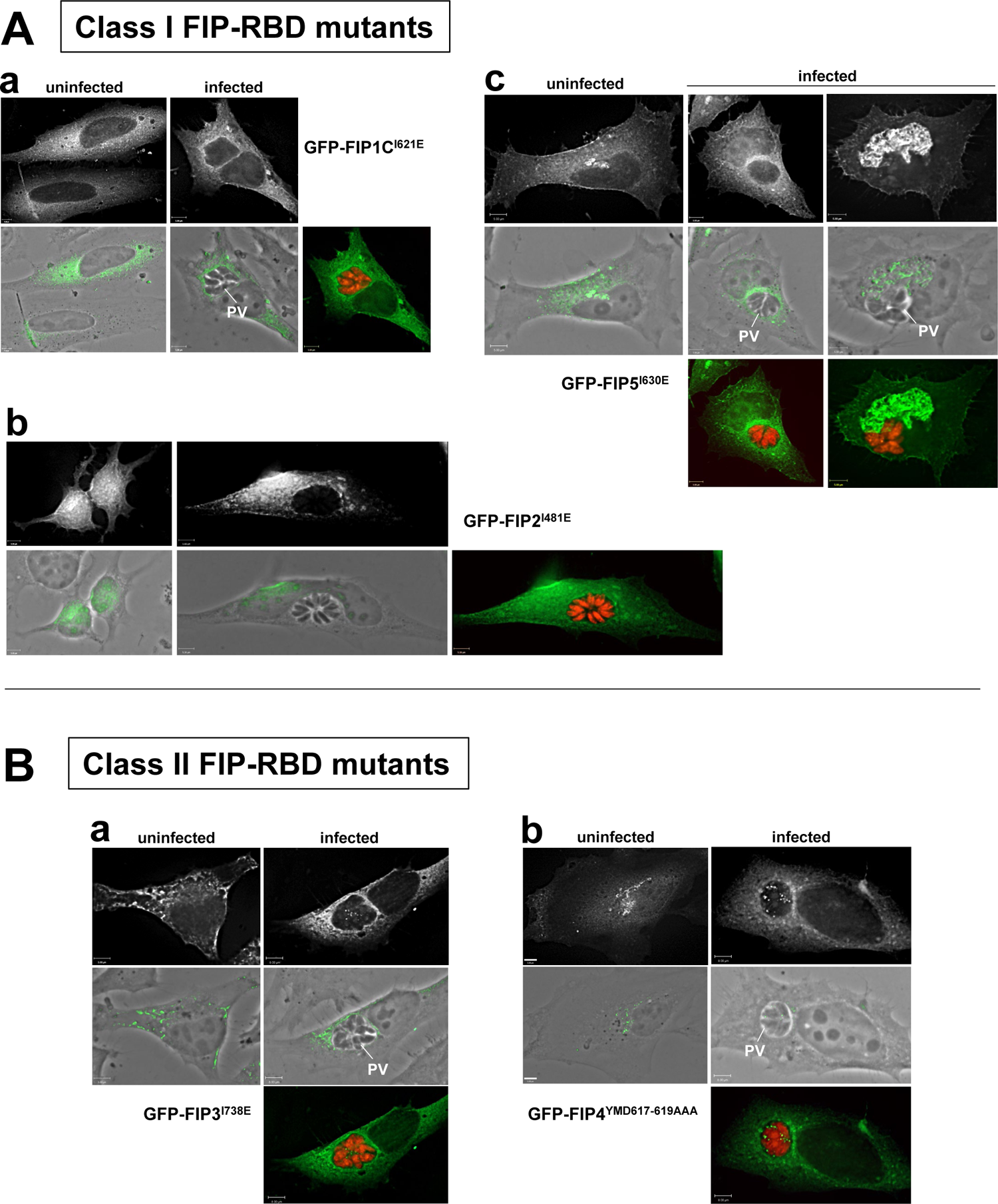
Distribution of mammalian vesicles containing FIP-RBD mutants from class I and II in *Toxoplasma*-infected cells. **A-B.** Fluorescence microscopy of HeLa cells transfected with FIP-RBD (Class I in A: GFP-FIP1C^I621E^, GFP-FIP2^I481E^ or GFP-FIP5^I630E^; Class II in B: GFP-FIP3^I738E^ or GFP-FIP4^YMD617-619AAA^), uninfected or infected with RFP-expressing *Toxoplasma* for 24h. Phase contrast overlaid with the GFP-FIP signal is shown as well as the GFP-FIP alone and with the RFP signal.

**Figure 5.**
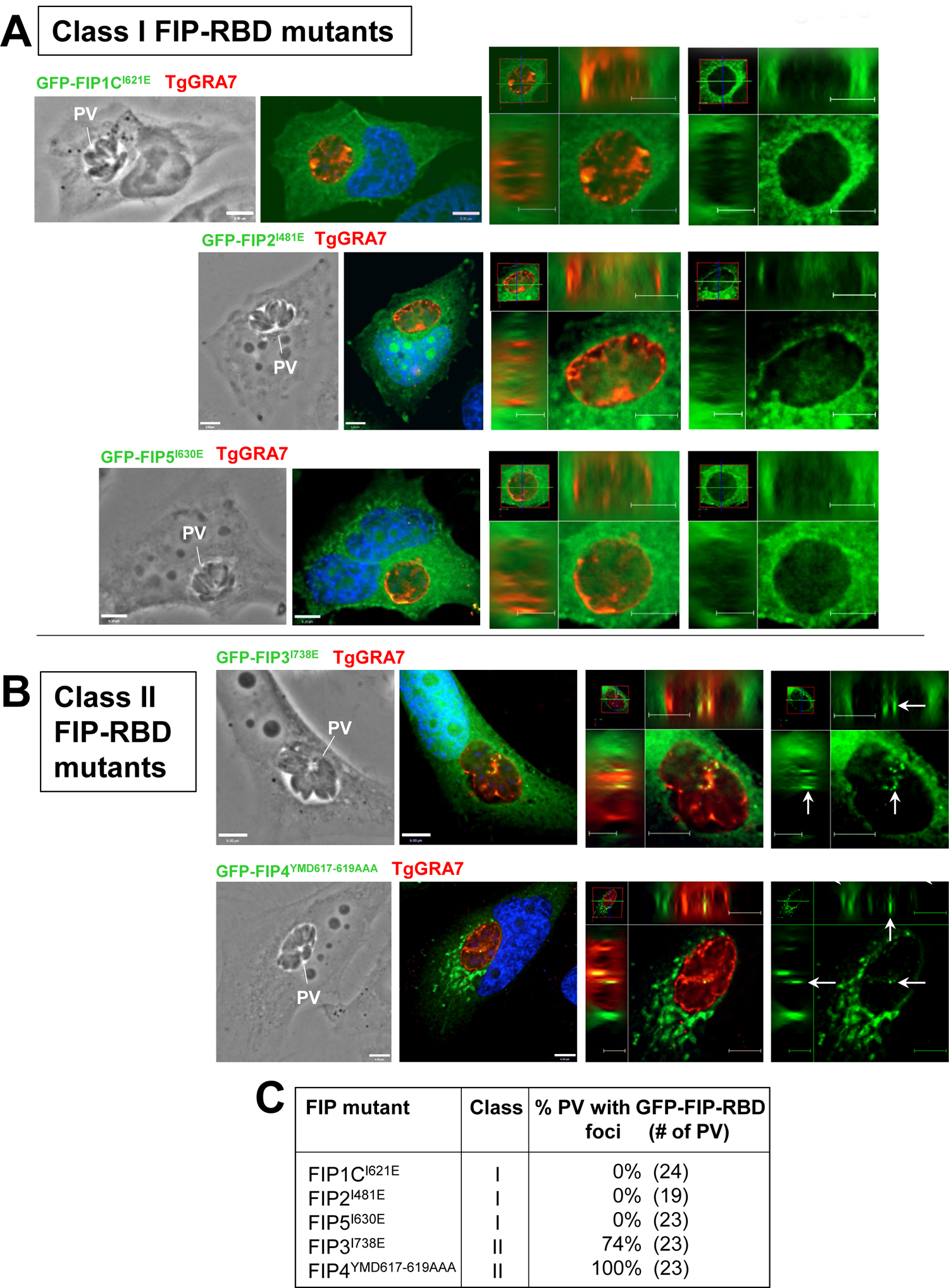
Distribution of mammalian vesicles containing FIP-RBD mutants from Class I and II in the *Toxoplasma* PV. **A-B.** Fluorescence microscopy of HeLa cells transfected with FIP-RBD mutants (Class I in A: GFP-FIP1C^I621E^, GFP-FIP2^I481E^ or GFP-FIP5^I630E^; Class II in B: GFP-FIP3^I738E^ or GFP-FIP4^YMD617-619AAA^), uninfected or infected with RFP-expressing *Toxoplasma* for 24h before immunostaining with anti-TgGRA7 antibody. DAPI in blue. For all images, individual z-slices and orthogonal views are shown. Arrows show intra-PV GFP-FIP foci. **C.** Quantification of the internalization of Class I and II FIP-RBD mutant vesicles within the *Toxoplasma* PV. Percent of PV with intra-PV foci was assessed based on PV viewed in A and B.

**Figure 6.**
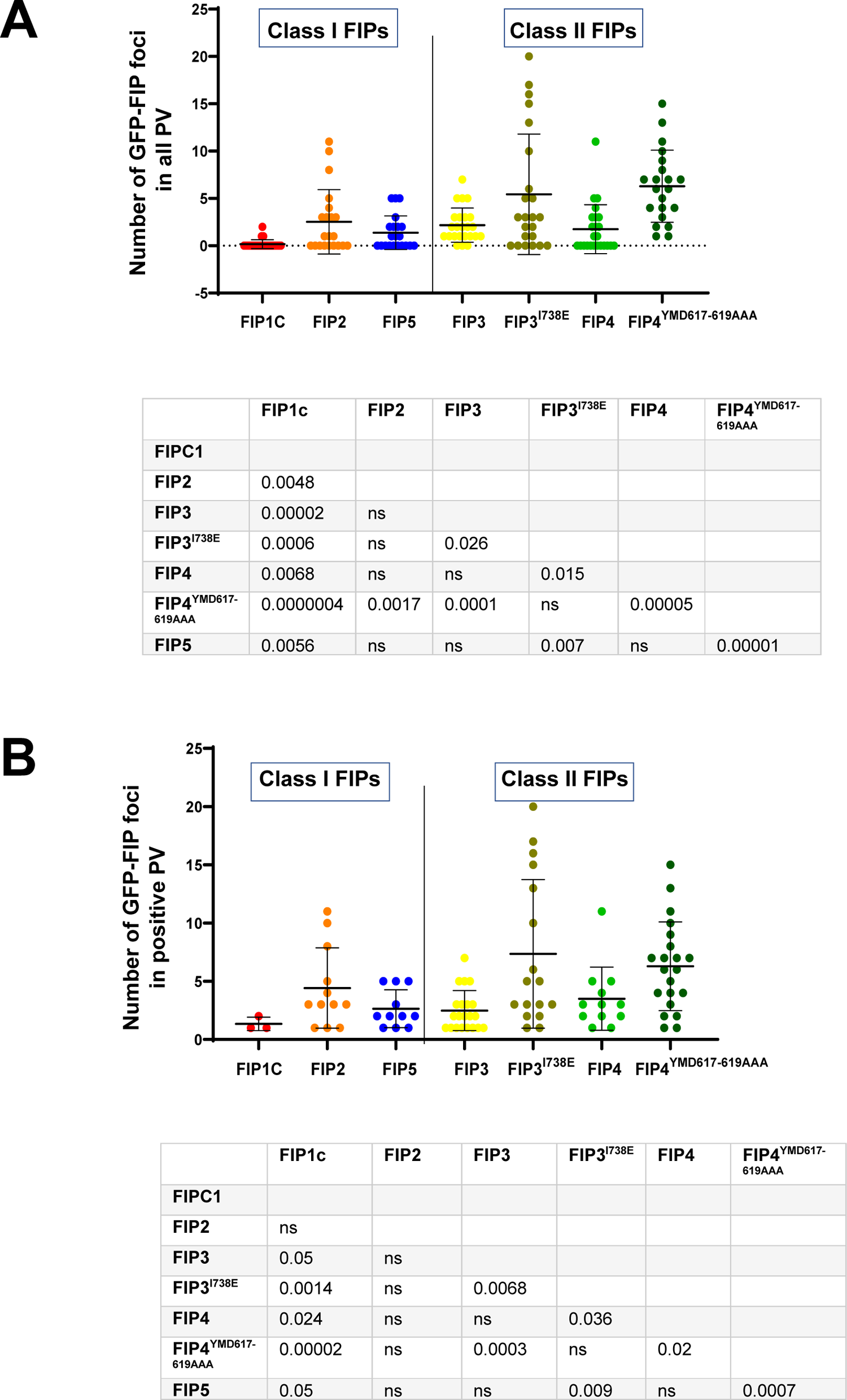
Enumeration of Rab11-FIP or FIP-RBD mutant vesicles within the *Toxoplasma* PV Intra-PV vesicles containing FIP or FIP-RBD mutants from class I and II were counted based on GFP foci in all the PV (A) or only in GFP-positive PV (B). Two-tailed *p* values shown. ns, *p* >0.05.

We conducted immunoEM studies on HeLa cells expressing GFP-FIP3 or GFP-FIP3-RBD mutant using anti-GFP antibody to scrutinize the GFP-positive vesicles around and inside the PV. In uninfected GFP-FIP3-expressing cells, gold particles were detected on the membranes of several vesicles and tubules in the cytoplasm (Figure 7A, panel a). In infected transfected cells, a noticeable gathering of labeled structures was observed around the PV (Figure 7A, panel b). Gold particles were also present on intra-PV vesicles (Figure 7A, panel c), and high magnification observations revealed the gold staining on vesicular membranes (Figure 7A, panel d). In HeLa cells expressing GFP-FIP3-RBD, the gold particle staining was more diffuse throughout the cell and partly associated with vesicles (Figure 7B, panel a). Similar to the GFP-FIP3 WT signal in infected cells, gold particles were observed in close proximity to the PV (Figure 7B, panel b); numerous labeled vesicles were observed in the PV lumen (Figure 7B, panel c), and membrane-associated on vesicles inside IVN tubules (Figure 7B, panel d).

**Figure 7.**
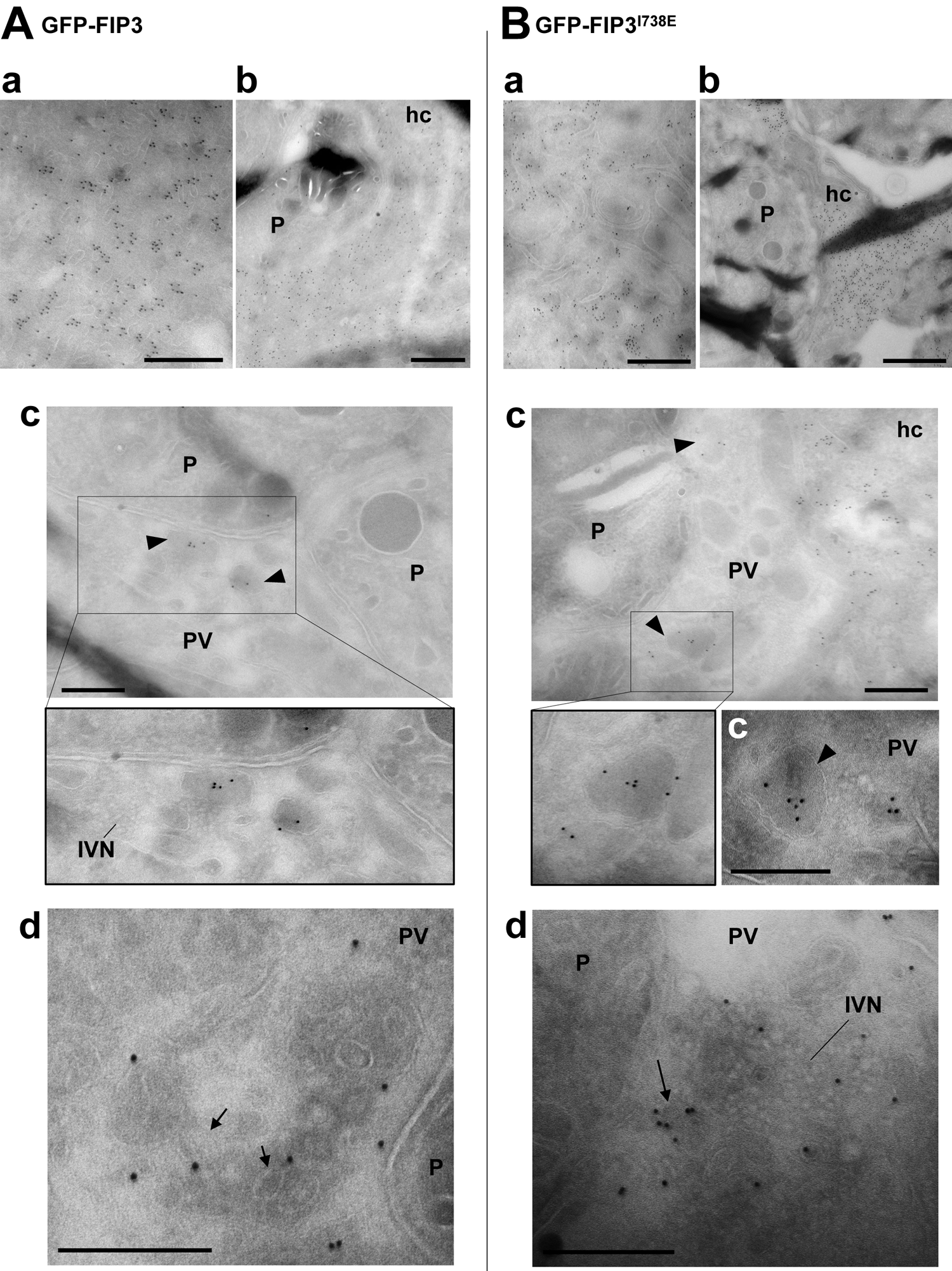
Immunolocalization of FIP3 and FIP3-RBD mutant in *Toxoplasma*-infected cells. **A-B.** ImmunoEM of HeLa cells expressing GFP-FIP3 (A) or GFP-FIP3^I738E^ (B) infected for 16h. FIP3 distribution was examined using anti-GFP antibody followed by protein A-gold particles. Panels a in A and B show gold particles in mammalian cells. Panels c and d in A and B illustrate gold particles localized within the PV on vesicles (arrowheads), and sometimes in the vicinity of the IVN. Some gold particles were also detected on the limiting membrane on internal vesicles (arrows). hc, host cell; P, parasite. All bars, 300 nm.

Jointly, these observations illustrate a striking difference between the two FIP classes regarding FIP vesicle internalization into the PV as the formation of Rab11-FIP complexes is required for Class I but is dispensable for Class II and even unfavorable based on the greater number of FIP3- and FIP4-RBD mutant vesicles detected in the PV.

### The PV contains mammalian vesicles marked with Rab11A, FIP3 or Rab11A-FIP3 complexes

The internalization of FIP3-positive vesicles into the PV independently of Rab11 led us to investigate the percentage of intra-PV Rab11A vesicles that are associated with FIP3. Transfected HeLa cells expressing GFP-FIP3 were immunolabeled with anti-Rab11A antibody to assess the colocalization of FIP3 and endogenous Rab11A proteins on vesicles. Expression of FIP3 in HeLa cells resulted in the congregation of Rab11A vesicles on pericentrosomal ERC with FIP3 (Figure 8A, panel a) as compared to non-transfected cells where Rab11A vesicles (detected by immunostaining) distributed throughout the cell with some concentration at the ERC (Figure 1A). The GFP-FIP3 signal was also distributed on ERC tubular extensions that contact Rab11A vesicles. In dividing cells, co-distributed GFP-FIP3 and Rab11A on vesicles were observed close to the intercellular cytoplasmic bridge formed between daughter cells during cytokinesis (Figure 8A, panel b), supporting targeted delivery of recycling endosomal Rab11 vesicles mediated by FIP3 to the midbody (Horgan *et al*., 2004; Wallace *et al*., 2002; Wilson *et al*., 2005; Fielding *et al*., 2005). Upon infection, the ERC, co-stained for GFP-FIP3 and Rab11A, was delocalized to the PV (Figure 8B, panel a). As illustrated in orthogonal views of *z*-slices, intra-PV vesicles marked for both FIP3 and Rab11A (Figure 8B, panel b), or only FIP3 or Rab11A were detected in close proximity to each other (Figure 8B, panel c) or distant (Figure 8B, panel d). Of 60 fluorescent foci analyzed from 14 PV, 27% of the foci were yellow (positive for both FIP3 and Rab11A), 50% red (Rab11A only) and 23% were GFP-labeled (FIP3 only). Although our assays could not take into account the presence of endogenous FIP3 associated with intra-PV vesicles, the detection of vesicles with GFP-FIP3 (excluding endogenous Rab11A) suggests the heterogenous composition of host recycling vesicles sequestered inyo the PV.

**Figure 8.**
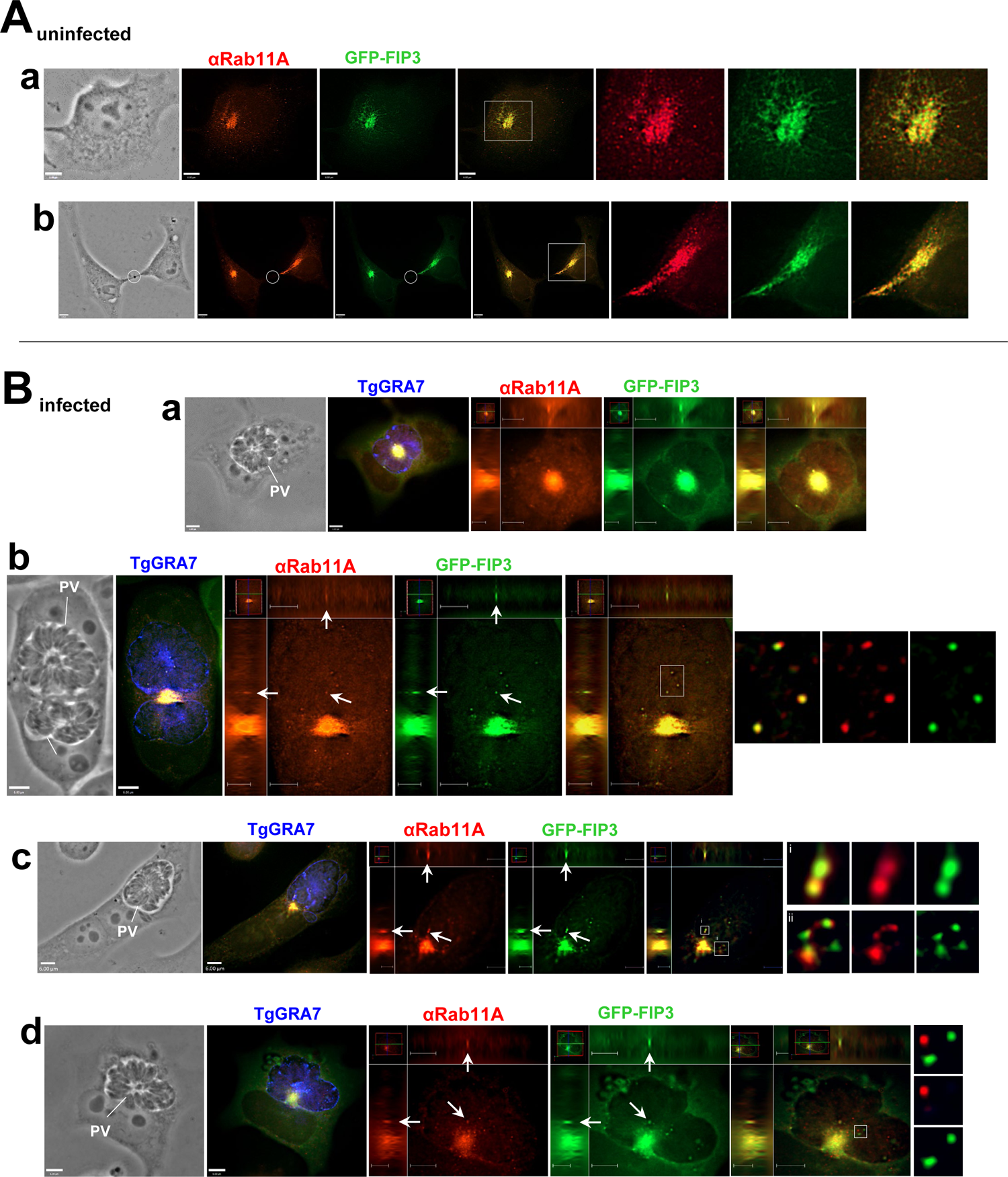
Abundance of mammalian Rab11A vesicles containing FIP3 in the PV. **A.** Fluorescence microscopy of HeLa cells transfected with GFP-FIP3 and immunostained with anti-Rab11A antibody at interphase (panel a) and during cytokinesis (panel b). The circle highlights the midbody. **B.** Fluorescence microscopy of HeLa cells transfected with GFP-FIP3 and infected for 24h before immunostaining for Rab11A and TgGRA7. The delocalization of the ERC at the top of the PV or squeezed between two PV is highlighted in panels a and b, respectively. For all images, individual z-slices and orthogonal views are shown. Arrows and squares in panels b-d show intra-PV GFP-FIP3 foci and/or Rab11A foci.

### Mammalian FIP3 vesicles are internalized in the PV in association with Arf6

The Rab11A-independent sequestration of FIP3 and FIP4 vesicles into the PV prompted us to investigate whether other FIP interactors were associated with Class II FIP vesicles in the PV lumen. Both FIP3 and FIP4 are Arf6 effectors (Shin *et al*., 2001). In mammalian cells, Arf6 localizes to the plasma membrane and endosomal compartments, and it is involved in the targeted delivery of recycling endosomal vesicles to the plasma membrane (D’Souza-Schorey and Chavrier, 2006). In preparation for cell division, Arf6 plays a vital role in targeting RE, e.g., Rab11 to the cleavage furrow and midbody (Schweitzer and D’Souza-Schorey, 2002). To compare the distribution of Arf6 in uninfected and *Toxoplasma*-infected cells, HeLa cells were transiently transfected with a plasmid containing Arf6 in fusion with C-terminal mCherry. In uninfected cells, the Arf6-mCherry signal was distributed in cytoplasmic structures with a higher concentration at the perinuclear region (Figure 9A). When transfected cells were infected for 24h, a strong perivacuolar staining was observed for Arf6-mCherry, suggesting interaction of the *Toxoplasma* PV with Arf6-positive vesicles (Figure 9B, panels a and b). Several Arf6-mCherry foci were observed within the PV, often concentrated at the IVN (Figure 9B, panel b).

**Figure 9.**
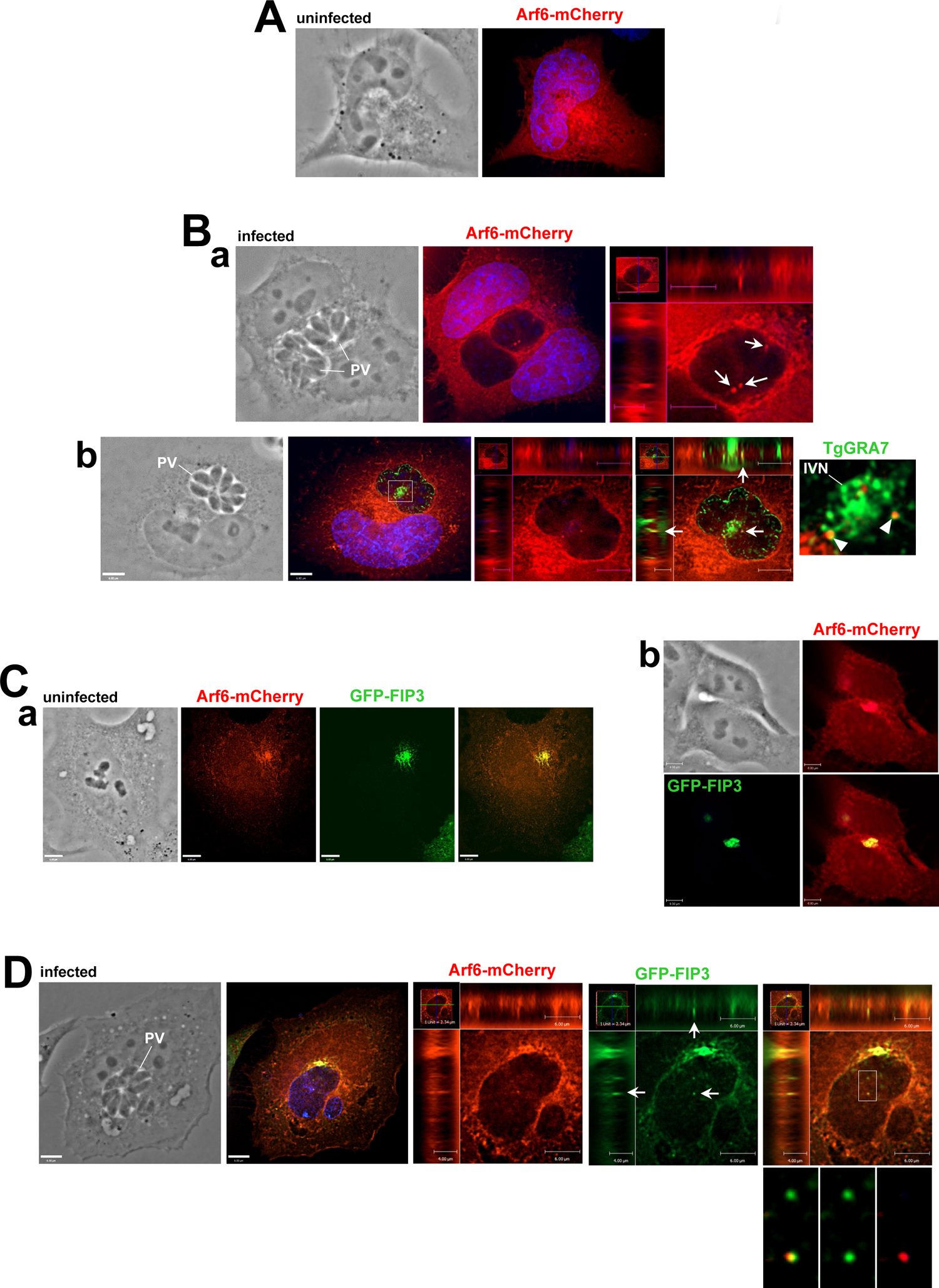
Distribution of mammalian vesicles containing Arf6 or FIP3 in *Toxoplasma*-infected cells. **A-B.** Fluorescence microscopy of HeLa cells transfected with Arf6-mCherry, uninfected (A) or infected for 24h (B), before immunostaining with anti-TgGRA7 antibody (panel b). For all images, individual z-slices and orthogonal views are shown. Arrows show intra-PV Arf6-mCherry foci (panel a) and arrowheads pinpoint foci associated with the IVN (panel b). **C-D.** Fluorescence microscopy of HeLa cells co-transfected with Arf6-mCherry and GFP-FIP3, uninfected (C, panel a: at interphase; panel b: end of cytokinesis) or infected for 24h (D). Arrows and squares show intra-PV Arf6-mCherry and/or GFP-FIP3 foci.

To examine whether the parasite recognizes vesicles with both Arf6 and FIP3, we transiently transfected HeLa cells with two plasmids, one encoding GFP-FIP3 and the other Arf6-mCherry. Previous studies reported that exogenously co-expressing FIP3 and Arf6 results in increased colocalization of the two proteins (Horgan *et al*., 2004; Fielding *et al*., 2005). In dually transfected cells, we confirmed an intense co-labeling of pericentrosomal ERC with GFP-FIP3 and Arf6-mCherry (Figure 9C, panel a). In dividing cells, the co-migration of vesicles positive for FIP3 and Arf6 was observed at the intersection between the two daughter cells, as illustrated in panel b in Figure 9C showing midbody remnants at the cell surface after abscission. In infected co-transfected cells, the Arf6-mCherry and GFP-FIP3 signals largely overlapped at the ERC close to the PV membrane and vesicles with Arf6 and FIP3 were observed within the PV (Figure 9D). In two independent assays, 86% and 61% of PV contained Arf6-mCherry vesicles upon mono-transfection or double transfection, respectively (Figure 10A). Approximately 70% of the Arf6-mCherry foci colocalized with the GFP-FIP3 foci were positive for Arf6-mCherry in cells expressing both Arf6 and FIP3. To determine whether Arf6 binding to FIP3 facilitates the Rab11-independent entry of class II FIP vesicles into the PV, we co-transfected HeLa cells with plasmids encoding Arf6-mCherry and GFP-FIP3-RBD mutant. The Arf6-mCherry and GFP-FIP3-RBD signals were dispersed in uninfected and infected cell, with partial colocalization (Figure 10B). The perivacuolar staining for Arf6-mCherry and GFP-FIP3-RBD was weaker in dually transfected and infected cells (Figure 10B) than for infected HeLa cells transfected with Arf6-mCherry alone (Figure 9B), GFP-FIP3 alone (Figure 2B, panel a), or Arf6-mCherry and GFP-FIP3 (Figure 9D). In these two independent assays, only ∼23% of PV contained Arf6-mCherry foci, and less than 30% of the Arf6-mCherry foci colocalized with the GFP-FIP3-RBD foci. There were also significantly less Arf6-mCherry foci per PV when co-transfected with GFP-FIP3-RBD as compared to co-transfection with GFP-FIP3 (Figure 11A). In comparison to the number of Arf6-mCherry foci per PV, the number of GFP-FIP3 or GFP-FIP3-RBD number of foci per PV remained constant upon double transfection. In fact, the number of GFP-FIP3 foci per PV increased following co-transfection with Arf6-mCherry while the number of Arf6-mCherry foci per PV dropped (Figure 11B).

**Figure 10.**
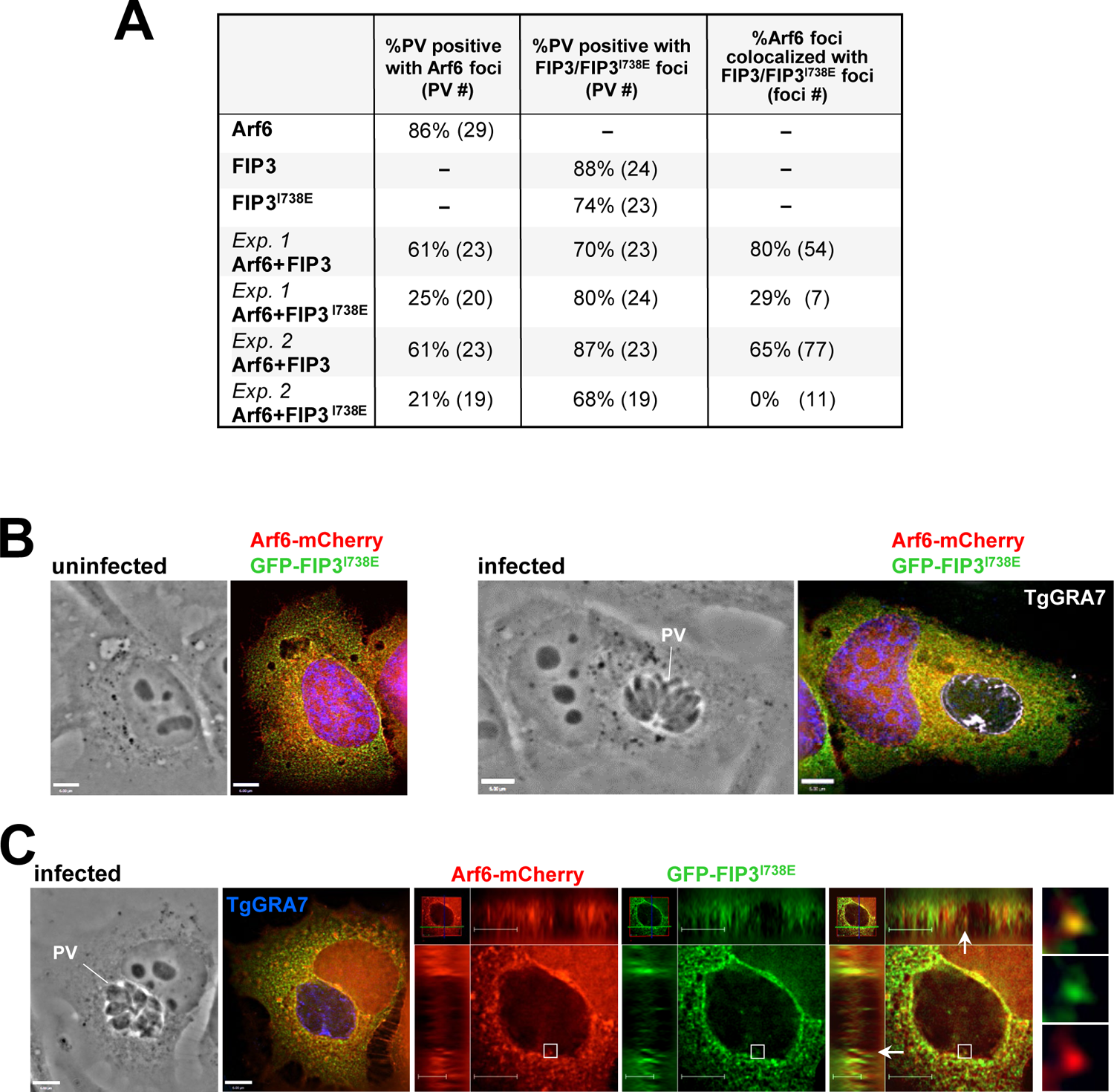
Distribution of mammalian vesicles containing Arf6 or FIP3-RBD in *Toxoplasma*-infected cells. **A.** Quantitative analysis of percent of PV containing Arf6-mCherry or GFP-FIP3, and the percent of Arf6-mCherry foci that colocalized with GFP-FIP3/GFP-FIP3-RBD based on experiments described in Figure 9 and below. **B-C.** Fluorescence microscopy of HeLa cells co-transfected with Arf6-mCherry and GFP-FIP3^I738E^, uninfected or infected for 24h, before immunostaining with anti-TgGRA7 antibody. For all images, individual z-slices and orthogonal views are shown. Arrows and squares in panels b-d show intra-PV GFP-FIP3-RBD foci.

**Figure 11.**
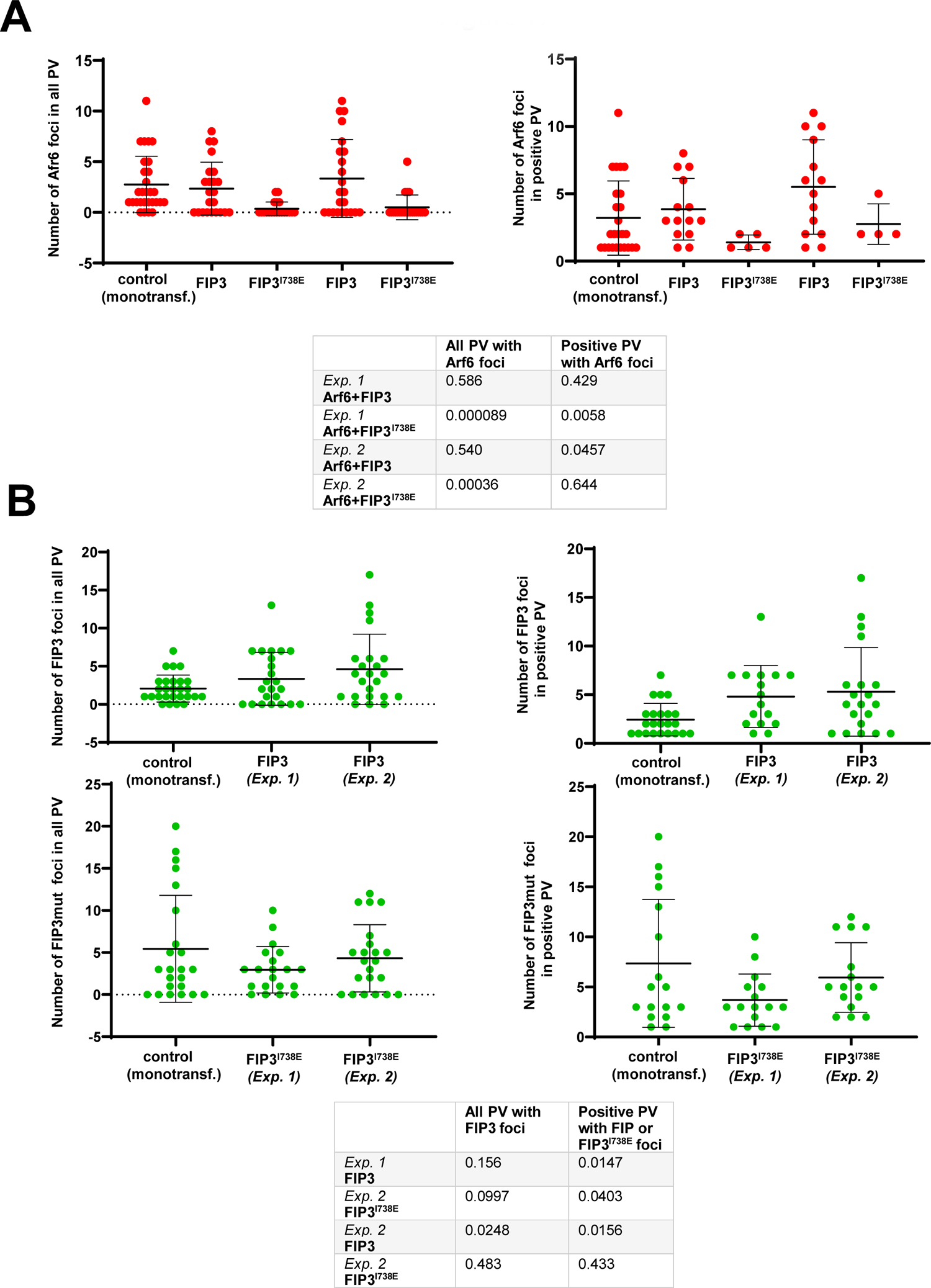
Enumeration of foci with Arf6, FIP3 or FIP3-RBD vesicles within the *Toxoplasma* PV. **A-B.** Quantification of data described in Figures 9 and 10. Intra-PV vesicles were counted in all the PV or all the positive PV for Arf6-mCherry (in A) or GFP-FIP3/GFP-FIP3-RBD (in B). Tables show two-tailed *p* values compared to control conditions corresponding to mono-transfected HeLa cells with Arf6-mCherry, GFP-FIP3 or GFP-FIP3^Δ687-712^.

These data suggest that the presence of Rab11 complexed to FIP3 leads to more efficient internalization of Arf6 vesicles into the PV and that Arf6-FIP3 interactions may not be the driving force for the delivery of vesicles of class II FIP-RBD mutants to the PV.

### Binding of FIP3 to either Rab11 or Arf6 increases the delivery of mammalian vesicles displaying FIP3 into the PV

FIP3 binds to the C-terminal α-helix of Arf6 (Schonteich *et al*., 2007). We generated a GFP-FIP3 construct with an altered ABD through the truncation of residues 687 to 712 to investigate further the relevance of FIP3 complexed to Arf6 for recognition and internalization of vesicles into the PV. HeLa cells were transfected with a plasmid encoding GFP-FIP3-ABD (FIP3^Δ687-712^) or a plasmid encoding the dual mutant GFP-FIP3-RBD-ABD (FIP3^I738EΔ687-712^), infected and analyzed by fluorescence microscopy. Numerous foci for GFP-FIP3-ABD vesicles were observed in the PV interior (Figure 12A) while no GFP-FIP3-RBD-ABD vesicles were detected in the vast majority of PV (Figure 12B, panel a). For PV in which GFP-FIP3-RBD-ABD foci were observable, the GFP signal was very weak and the foci were tiny (Figure 12B, panel b) compared to foci for GFP-FIP3 WT or GFP-FIP3-RBD. The percentage of PV containing vesicles with GFP-FIP3-ABD and the number of those vesicles as quantified in Figure 12C were comparable to the percentage of PV and number of vesicles for GFP-FIP3 WT or GFP-FIP3-RBD vesicles (Figures 3C, 5C and 6). However, only 37% of PV were positive for the dual mutant GFP-FIP3-RBD-ABD, and the intra-PV foci for GFP-FIP3-RBD-ABD were very tiny.

**Figure 12.**
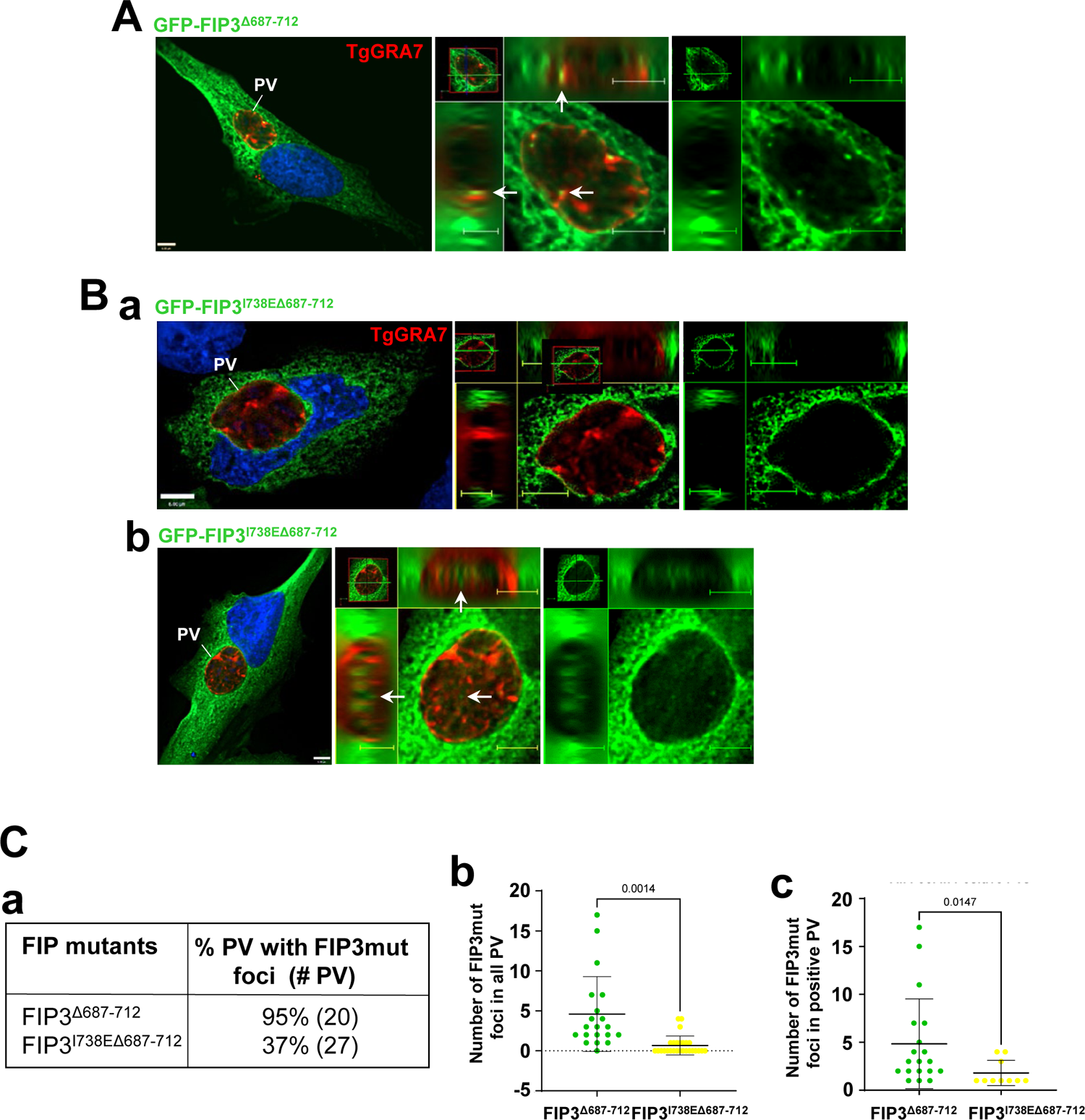
Comparative distribution of mammalian vesicles containing Rab11-FIP3-ABD and Rab11-FIP3-RBD-ABD in the *Toxoplasma* PV. **A-B.** Fluorescence microscopy of HeLa cells co-transfected with GFP-FIP3^Δ687-712^ (A) or GFP-FIP3^I738EΔ687-712^ (B, panels b and c) and infected for 24h before immunostaining with anti-TgGRA7 antibody. DAPI in blue. For all images, individual z-slices and orthogonal views are shown. Arrows show intra-PV foci. **C.** Percent of PV with intra-PV foci corresponding to vesicles with GFP-FIP3-ABD or GFP-FIP3-RBD-ABD (panel a) and enumeration of mammalian intra-PV vesicles with GFP-FIP3-ABD or GFP-FIP3-RBD-ABD mutants in all PV (panel b) or PV positive for GFP (panel c). Two-tailed p-values are shown.

These data suggest that Rab11-FIP3 or Arf6-FIP3 interactions may not be individually require to drive FIP3 vesicle sequestration into the PV but abrogating the binding of both Rab11 and Arf6 to FIP3 is detrimental for this process. However, since abrogation of both binding processes does not completely impede vesicle delivery to the PV, other FIP3 interactors may drive the internalization. A graphical summary of our data presented in Figure 13 illustrate the Class I FIP dependence on Rab11 binding for internalization and the PV’s ability to internalize FIP3 vesicles depending on whether their interaction with Rab11, Arf6, or neither.

**Figure 13.**
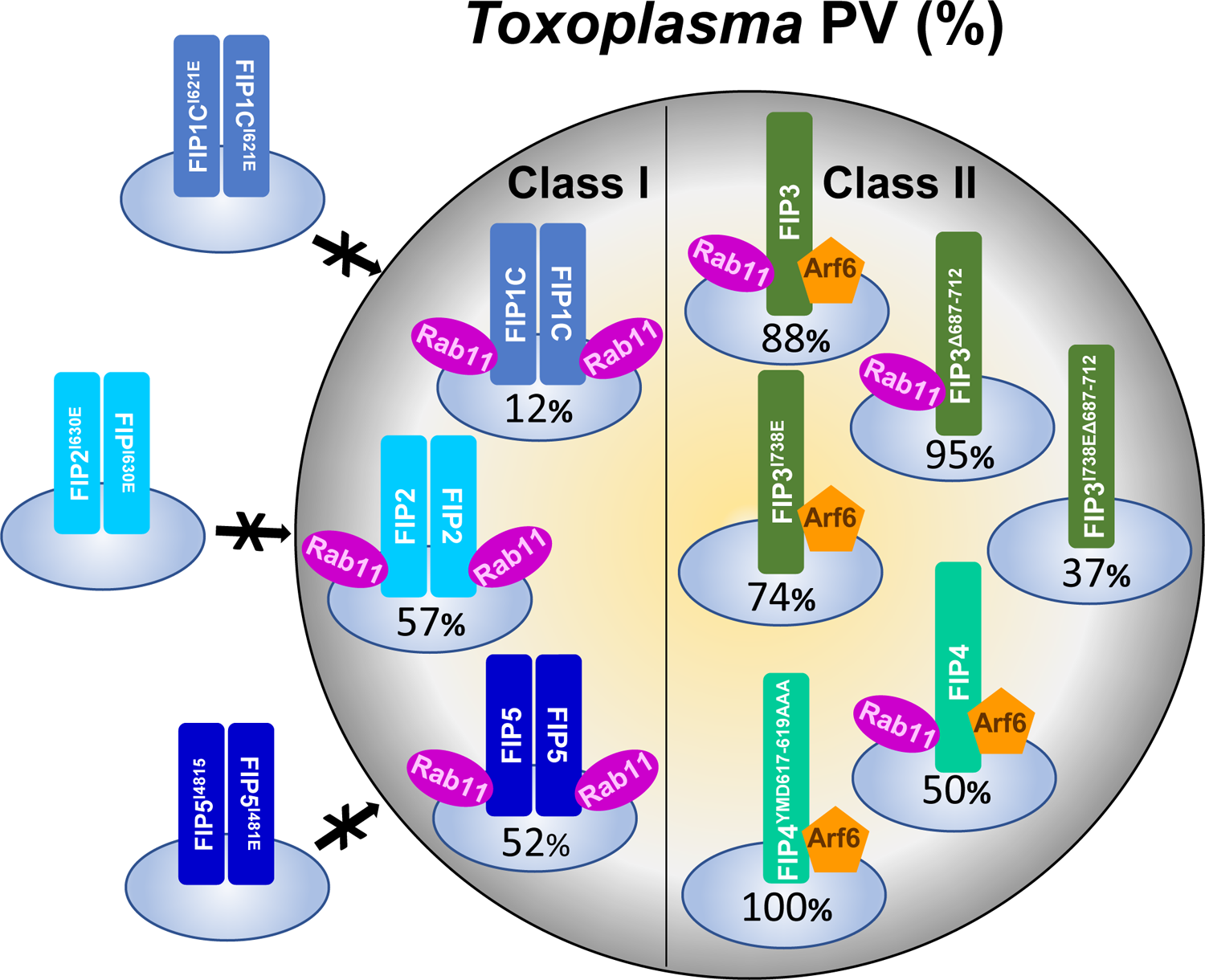
Graphical summary of the differential internalization of mammalian FIP recycling vesicles into the *Toxoplasma* PV The percentage of positive PV contain FIP-associated vesicles is shown. The percentage of PV containing recycling vesicles marked with a WT or RBD-mutant FIP from Class I or Class II is indicated, revealing selectivity for PV internalization among FIP vesicles. No vesicles with Class I FIP-RBD mutants are detected in the PV while vesicles with class II FIP-RBD mutants are still internalized, with PV contain vesicles with FIP4-RBD. Vesicles with FIP3-RBD or FIP3-ABD are internalized into PV. Very few PV display vesicles with the dual mutant FIP3-RBD-ABD.

### *Toxoplasma* infection results in the altered expression of several genes involved in the host mammalian recycling system

The intersection of the *Toxoplasma* PV with mammalian Rab11, FIPs and Arf6 proteins that regulate recycling endocytosis or cytokinesis led us to examine more broadly the impact of infection on the host recycling endosomal network. We performed RNA sequencing analysis on *Toxoplasma*-infected fibroblasts 24h post-infection to survey alterations in the expression of genes encoding Rabs, Rab GTPase activating proteins (RabGAP) and Rab guanine nucleotide exchange factors (RabGEF), which control the recycling system (Figure 14; see details of all genes in Table SI and description some effectors in Figure S3). Among Rab proteins, Rab15, which co-localizes with Rab4, Rab5 and the TfR on sorting endosomes (SE) as well as with Rab11 on the ERC (Zuk and Elferink, 1999; 2000), is up-regulated upon infection. Rab15 regulates the TfR trafficking through SE and the ERC, and influences iron levels in mammalian cells. Concerning Rab5 proteins, the Rab5b isoform, which localizes to a TfR-positive endosomal compartment (Wilson and Wilson, 1992; Bucci *et al*., 1995), is down-regulated in infected cells. Upon *Toxoplasma* infection, no changes in expression levels were detected for other Rabs involved in recycling such as Rab11 and Rab35; Rab35 is a major player in the fast recycling pathway and cytokinesis and localizes at the plasma membrane and endosomes (Klinkert and Echard, 2016).

**Figure 14.**
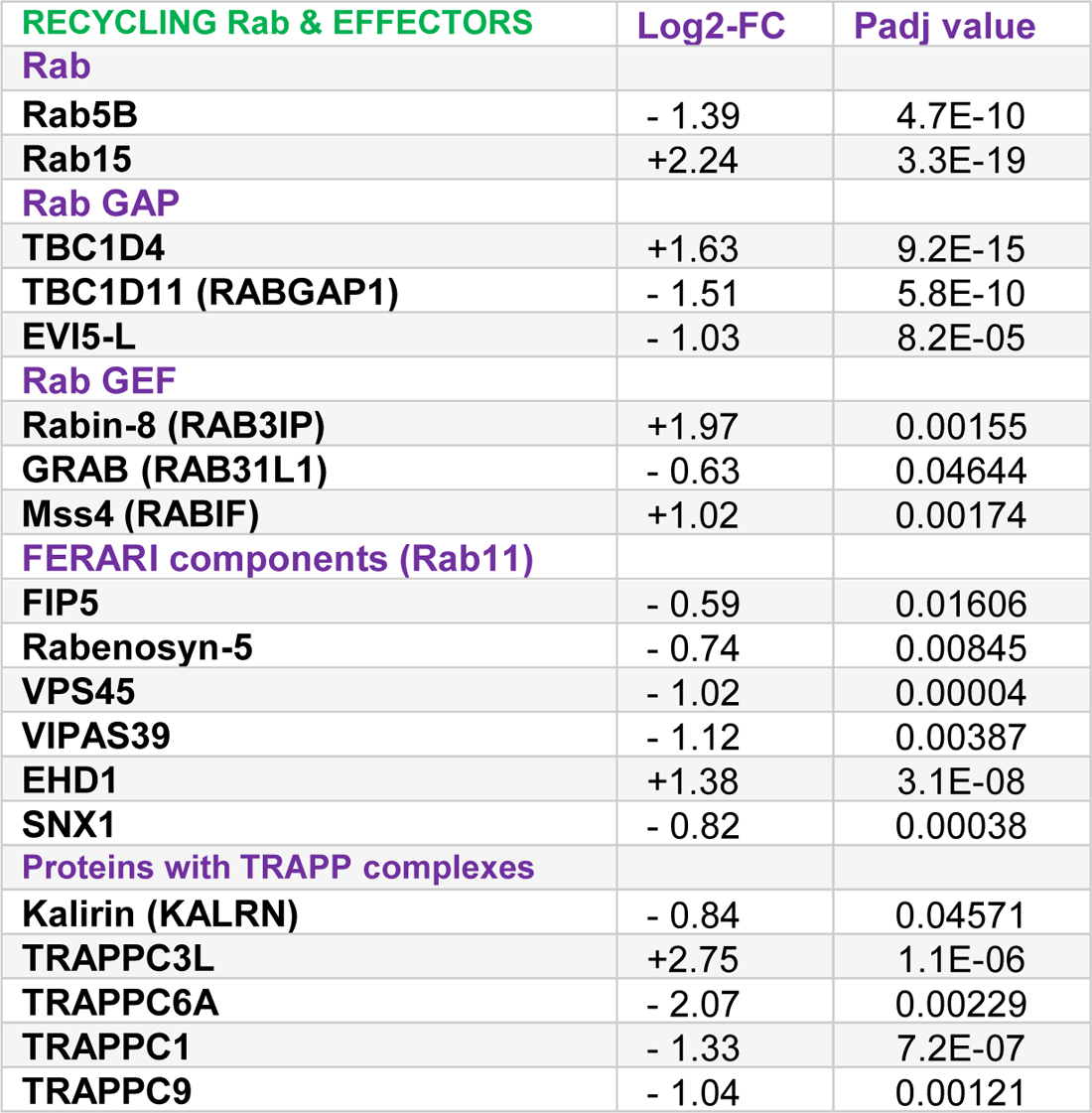
Changes in gene expression for host recycling Rab proteins and effectors upon a 24h *Toxoplasma* infection in fibroblasts RNA-seq data showing Log_2_ fold change in gene transcripts identified in the database and padj values of statistical significance.

TBC domain-containing proteins (TBC proteins) function as RabGAP, promoting the hydrolysis of Rab-GTP to Rab-GDP for the regulation of specific intracellular trafficking pathways. TBC1D4 enhances glucose uptake in cells through glucose transporter 4 (GLUT4) (Kane *et al*., 2020; Miinea *et al*., 2005; Sano *et al*., 2003; Treeback et al., 2010) and is up-regulated in infected cells. TBC1D11, which is down-regulated in infected cells, is involved in cytokinesis through Arf6-positive RE recruitment (Kanno *et al*., 2010; Kobayashi *et al*., 2014). The Rab11 GAP EVI5 is also down-regulated in infected cells, is a modulator of cell cycle progression (Lim and Tang, 2013), and co-localizes with TBC1D11 in RE containing TfR (Kanno *et al*., 2010).

After prenylation, Rab proteins are delivered to a target membrane where they are activated by a GEF to replace bound GDP by GTP. The Rab8 GEF Rabin-8 is up-regulated in infected cells and it controls the exocytic trafficking of proteins from the TGN and RE vesicles containing TfR to the plasma membrane (Feng *et al*., 2015) and co-distributes with Arf6 for vesicle delivery to the plasma membrane (Hattula *et al*., 2006; Peranen, 2011; Feng *et al*., 2015). GRAB (RAB31L1) is down-regulated in infected cells and is implicated in the exo-endocytic cycling of vesicles (Pavlos *et al*., 2010). Mss4 (RabIF) is up-regulated upon infection, and this holdase chaperone promotes the stability of Rab10, a key Rab in GLUT4 exocytosis (Gulbranson *et al*., 2017).

Factors for Endosome Recycling And Rab Interactions (FERARI) is a tethering platform for multiple effectors acting in Rab11 recycling pathways at SE (Solinger *et al*., 2020). The Rab-binding module of FERARI consists of FIP5 (binds to Rab11) and rabenosyn-5 (binds to Rab4 and Rab5), while the SNARE-interacting module comprises VPS45 and VIPAS39 (de Renzis *et al*., 2002; Solinger *et al*., 2020; van der Kant *et al*., 2015; Mohrmann *et al*., 2002). All four of these components from FERARI are down-regulated in infected cells. Endocytosed transferrin is sequentially transported through Rab5, Rab4 and Rab11 vesicles before recycling back to the plasma membrane (Sonnichsen *et al*., 2000; Ullrich *et al*., 1996; Daro *et al*., 1996). Rabenosyn-5 stimulates TfR recycling by activating the fast pathway and switching off the slow recycling route through Rab11-positive membranes (Mohrmann *et al*., 2002). FERARI, which also promotes membrane fission and fusion of SE at the ERC, includes the dynamin-like protein EHD1 (Daumke *et al*., 2007) that is up-regulated upon *Toxoplasma* infection. EHD1 exerts its pinching activity on tubular endosomal membranes, playing a role in ERC maintenance (Solinger *et al*., 2020).

The TRAnsport Protein Particle (TRAPP) is a modular multiunit complex that acts as a GEF for Rab proteins and thus is an important regulator of membrane traffic (Lipatova *et al*., 2019; Sacher *et al*., 2019). Upon infection, four TRAPP components have dysregulated expression, including TRAPPC1, TRAPPC6A, and TRAPPC3L that are part of the TRAPP core (Kim *et al*., 2016). Kalirin, which is one of the four subunits forming a core that confers the GEF activity onto TRAPP, forms a complex with several TRAPP, including TRAPPC9 to facilitate the recycling activities of Rab11 such as for TfR (Kim *et al*., 2006; Wang *et al*., 2020). Of the 5 TRAPP components with dysregulated expression, 4 out 5 are down-regulated, which suggests decreased preference for the slow Rab11 recycling of TfR.

Overall, several genes encoding effectors involved in recycling pathways (e.g., for GLUT4 and TfR) have differentiated expression in *Toxoplasma*-infected cells compared to uninfected cells, suggesting the manipulation of many host recycling compartments and effectors by the parasite, perhaps to increase nutrient (e.g., glucose, iron) pools in the host cell.

## Discussion

In this study, we reported that the intravacuolar parasite *Toxoplasma* manipulates the trafficking pathways of mammalian recycling endosomes (RE) that contain lipids as cargoes destined for the plasma membrane for re-use by the cell or for membrane donation to form the cleavage furrow during cytokinesis. Mammalian Rab11 proteins regulate the traffic of the slow recycling pathway for many cell surface proteins, including TfR, ion channels, junctional proteins, integrins, and immune receptors (Campa & Hirsch, 2017). We have shown that Rab11A and Rab11B vesicles distribute around the PV and are internalized into the vacuole (Romano *et al*., 2017), and that subsets of Rab11 vesicles with Class I and class II FIP are differentially scavenged by *Toxoplasma* (this study). A previous study showed that ectopically expressed Arf6 surrounds the PV in infected VERO cells during invasion, which potentially facilitates PV membrane biogenesis (da Silva *et al*., 2009). Here, we demonstrated that the perivacuolar localization of mammalian Arf6 persists throughout infection and that Arf6 vesicles penetrate the PV. Finally, we observed that the ERC is dramatically delocalized from its pericentrosomal region of the host cell to be associated with the PV, underlying important alterations in the recycling system in the host cell.

*Why does Toxoplasma divert host Class I FIP vesicles?* Class I FIPs function in endosomal recycling processes (reviewed in Jing and Prekeris, 2009) and bind to Rab11 to govern the trafficking of Rab11 vesicles to the cell surface. The C2-domain of class I FIPs targets the Rab11-FIP complexes to docking sites on the plasma membrane selectively enriched in phosphatidylinositol-3,4,5-trisphosphate and phosphatidic acid, two lipids required for fusion (Lindsay and McCaffrey, 2004b). *Toxoplasma* may advantageously intercept Class I FIP vesicles to scavenge their cargo for nutritional benefit and/or to neutralize toxic host factors through their sequestration into the PV. Among specific cargo trafficked to the cell surface by Class I FIP, the glucose transporter GLUT4 is trafficked to the cell surface via FIP5, which partially localizes to intracellular GLUT4 storage vesicles (GSV) (Welsh *et al*., 2007). FIP1C mediates the recycling of several plasma membrane receptors, including TfR, and FIP2 is implicated in the recycling of many molecules, e.g., GLUT4, TfR, the water channel protein aquaporin-2 and the chemokine receptor CXCR2 (Horgan and McCaffrey, 2009).

The cargo of Class I FIP vesicles may supply nutrients for replicating *Toxoplasma*. For example, the source of iron for *Toxoplasma* is still unknown, and FIP1 and FIP2 vesicles associated with TfR-Tf may provide iron to the parasite. Of interest, our RNA seq data reveal that genes coding for regulators of TfR recycling, such as TBC1D11, EVI5, Rabenosyn-5 and Kalirin, have altered expression upon infection and these changes may affect the TfR cycle leading to the retention of iron-loaded transferrin in infected cells to the parasite’s profit. GLUT4 is the main glucose transporter in muscle and fat cells, and it localizes at the plasma membrane and many organelles such as SE, RE and the TGN as well as GSV that mediate the transport of GLUT4 between these compartments (Bryant *et al*., 2002). In the absence of insulin, about ∼35% of GLUT4 remain trapped in an intracellular circuit between endosomes and the TGN (Martin *at al*., 1996). In fibroblasts (used in our RNAseq screen), most GLUT4 localizes with TfR in endosomes (Lampson *et al*., 2001). TBC1D4 that shuttles GLUT4 between GSV and the plasma membrane, is up-regulated in *Toxoplasma*-infected cells, potentially leading to increased glucose intake, which may be beneficial for the parasite. Moreover, the RabGEF Mss4 that stabilizes Rab10, a GTPase involved in GLUT4 exocytosis and, thus mediating GLUT4 surface-exposure, is up-regulated upon infection. In skeletal muscle cells, GLUT4 is highly expressed to promote glucose intake in response to insulin for energy demands. The tropism and attraction of *Toxoplasma* for skeletal muscle cells for encystation (Swierzy and Lüder, 2015) may be linked to the metabolic properties of skeletal muscle cells, such as a high glucose supply, which represent a propitious environment for the long-term survival of the parasitic chronic stage.

*Toxoplasma* may also target Class I FIP vesicles to modulate the immune response of its host. For example, the chemokine receptor CXCR2 is present into Rab11A-positive RE (Fan *et al*., 2003), and its recycling and thus receptor-mediated chemotaxis is regulated by FIP2 (Fan *et al*., 2004). CXCR2 stimulates cell migration in response to a concentration gradient of the CXC chemokine IL-8 ligand (Wolf *et al*., 1998), and is essential for the recruitment of leukocytes to inflammatory sites (Garcia-Ramallo *et al*., 2002). Neutrophils employing a battery of microbicidal activities, e.g., phagocytosis, neutrophil extracellular trap (NET) formation and release of peroxide species, are required for early resistance against *Toxoplasma* in the gut (Sayles and Johnson, 1997; Bliss *et al*., 2001). In particular, CXCR2 is involved in early neutrophil recruitment and plays an important protective role in resistance to *Toxoplasma* (Del Rio *et al*., 2001). In fact, *Toxoplasma* infection of CXCR2^−/−^ mice results in defects in neutrophil migration to the site of infection, lower production of proinflammatory cytokines (e.g., TNF-α, IFN-γ), and higher brain cyst numbers during chronic infection, compared to WT mice. To this point, the sequestration of vesicles containing CXCR2-FIP2 complexes into the PV may be advantageous for the parasite to avoid neutrophil attraction to infected tissues, thus ensuring parasite dissemination in the host.

In addition to CXCR2, the cargoes of Rab11 vesicles include immune mediators (Kelly *et al*., 2012): TNF-α, which triggers IFN-γ-primed macrophages for *Toxoplasma* killing (Sibley *et al*., 1991); IL-6, which protects during *Toxoplasma* early infection by promoting IL-17 production from NK cells for T-cell activation (Jebbari *et al*., 1998; Passos *et al*., 2010) and TLR4, which is a critical innate immune cell receptor involved in *Toxoplasma* detection and activation of immune responses (Zare-Bidaki *et al*., 2014). Diversion of these anti-*Toxoplasma* effectors to the PV could be part of a strategy crafted by the parasite to thwart host cell-intrinsic innate immunity pathways.

The internalization of Class I FIP vesicles is Rab11-dependent, suggesting the recognition of complexes formed between FIP1, FIP2 or FIP5 and Rab11. This would ensure an elective and efficient targeting of Class I FIP-Rab11 RE by the parasite to benefit from their cargoes.

*Why does Toxoplasma divert host Class II FIP vesicles?* While class I FIP are specifically implicated in the regulation of endocytic protein sorting and recycling, class II FIP have different roles and more specialized functions depending on the cell cycle (reviewed in Jing and Prekeris, 2009). FIP3 and FIP4 localize to the ERC and act as scaffold proteins for this organelle, tethering the ERC to the pericentrosomal region (in many cell types) through binding to microtubule motor proteins. Knock-down of FIP3, overexpression of FIP4 or alteration of the microtubule cytoskeleton (e.g., with nocodazole) causes abnormal distribution of the ERC to the cell periphery (Horgan *et al*., 2007; Wallace *et al*., 2002; Lin *et al*., 2002). In *Toxoplasma*-infected cells, the host ERC is dragged toward the PV. This process may be consecutive to the hijacking of the host MTOC by the parasite, resulting in the rearrangement of the host microtubular network all around the vacuole (Coppens et al., 2006; Walker et al., 2008). Manipulating the host microtubular network allows for the parasite to govern the host cellular membrane transport and attract organelles to the PV. Of importance, the recruitment of the host MTOC and microtubules is actively mediated by the parasite as these events also occur in mammalian cytoplasts that lack genetic materials and are, in other words, dying fragments of cytoplasm (Romano et al,. 2008). The co-option of host microtubules is also associated with supranumerous foci of γ-tubulin and aberrant multi-nucleated structures in infected cells, leading to cell cycle dysregulation and defects in cytokinetic events. By stalling host cellular division and blocking cytokinesis, the parasite may insure a vast and favorable environment for replication by increasing resource availability, such as lipid membranes that would normally be targeted to the cleavage furrow to complete cytokinesis. During cytokinesis, FIP3 or FIP4 associated with Rab11 moves Rab11 RE to the cleavage furrow via centrosome-anchored microtubules (Fielding *et al*., 2005). At the furrow, the Rab11-FIP3 or Rab11-FIP4 vesicles encounter and interact with Arf6 on the plasma membrane, resulting in tethering of these vesicles to the plasma membrane via interaction with the Exocyst complex. In this case, the attraction and retention of FIP3, FIP4 and Arf6 vesicles in the PV may represent an additional stratagem developed by *Toxoplasma* to cause cytokinesis failure by blocking the abscission process.

Alternatively, the primary target of *Toxoplasma* may be the host ERC, which is pulled toward the PV subsequently to MTOC capture. The ERC is a long-lived compartment, with multiple sorting functions toward the recycling, retrograde and exocytic routes (Maxfield and McGraw, 2004). Most proteins that cycle between the cell interior and the cell surface accumulate in the ERC because this is the slow step in their return to the plasma membrane. For example, all of the LDL receptors and TfR recycle from the ERC to the cell surface. Rab15 regulates iron levels in the ERC (Zuk and Elferink, 1999; 2000); Rab15 up-regulation in infected cells may result in an increase in iron content in the ERC. The ERC is also a major repository of cholesterol that enters the organelle by endocytosis and non-vesicular mechanisms (Hao *et al*, 2002). A prime example of essential metabolites for the parasite is cholesterol, which is derived from LDL that are internalized into the host cell (Coppens *et al*., 2000). *Toxoplasma* also relies on host sphingolipids for optimal replication (Romano *et al*., 2013), and sphingomyelin transits through the ERC prior to delivery to various intracellular destinations (Koval and Pagano, 1989). The ERC may then represent a source of endogenous and exogenous metabolites for *Toxoplasma*, and targeting the ERC for its massive pool of cholesterol and sphingolipids would be advantageous for the parasite.

The proximity of the ERC to the PV would also facilitate the scavenging of FIP3 and FIP4 vesicles by the parasite. The ERC is composed of both Rab11- and Arf6-positive membrane subsets. The trapping of FIP3 and FIP4 vesicles inside the PV is Rab11-independent, suggesting a mechanism of internalization for class II FIP vesicles different from Class I FIP vesicles, such as the mediation of Arf6 for recognition and scavenging of class II FIP vesicles. Overall, our data highlight a selective process of interaction with host vesicular components at the PV membrane, likely mediated by specific parasite proteins.

Arf6 mediates a clathrin-independent endocytosis (CIE) pathway, in which cargo is internalized and transported in Arf6-enriched vesicles to be subsequently recycled to the plasma membrane (Van Acker *et al*., 2019). Thus, like Rab11-associated vesicles, Arf6-associated endosomes may play roles in nutrient acquisition or immune response control. Arf6-associated endosomes carry cholesterol and Arf6 is a central regulator of cholesterol homeostasis, controlling both the uptake and efflux of this lipid from cells (Naslavsky *et al*., 2004; Schweitzer *et al*., 2009). Therefore, the intra-PV Arf6 vesicles may constitute a providential source of cholesterol for *Toxoplasma*. Additional potential nutrient modulating cargo for Arf6-associated endosomes include the glucose transporter GLUT1 and some amino acid transporters. MHC class I molecules are also present on Arf6-positive tubules emanating from the ERC. It has been reported that cross-presentation by MHC class I molecules, which allow the detection of exogenous antigens by CD8^+^ T lymphocytes, is crucial to initiate cytotoxic immune responses against *Toxoplasma* (Blanchard *et al*., 2008). Interestingly, the Δgra2Δgra6 *Toxoplasma* mutant that lacks the IVN and is impaired in intra-PV vesicle internalization, is more susceptible to MHC I presentation than WT parasites (Lopez *et al*., 2015). This suggests that the scavenging of Arf6 vesicles could be an immune regulatory process mediated by the parasite to interfere with antigen cross-presentation and immune detection.

Like *Toxoplasma*, several intracellular bacteria target the mammalian host endocytic recycling circuitry (Brumell and Scidmore, 2007; Allgood and Neunuebel, 2018). By usurping host recycling Rab (e.g., Rab4, Rab11, Rab35), bacteria promote their entry and exit from the host cell, control the biogenesis, maturation or rupture of their vacuole, and evade immune detection. From a therapeutic point-of-view, any essential parasite and bacterial effectors located at the vacuolar membrane or secreted into the host cell for interaction with specific Rab proteins could represent potential drug targets to combat microbial infections. Seminal information on the role of mammalian host Rab effectors has come from work on intracellular pathogens that co-opt Rab functions (Stein *et al*., 2012; Sherwood and Roy, 2013; Damiani *et al*., 2014). From a cell biology perceptive, studying the unidirectional transit of host Rab vesicles toward the *Toxoplasma* PV may reveal which Rab effectors bind to parasite proteins at the PV, perhaps expanding our repertoire of Rab-interacting proteins.

## Materials and Methods

### Reagents and antibodies

All reagents were purchased from Sigma (St. Louis, MO) or Thermo Fisher Scientific (Waltham, MA), unless otherwise stated. The primary antibodies used in this study were: rat and rabbit polyclonal anti-GRA7 (Coppens *et al*., 2006), rabbit anti-Rab11 (Cell Signaling Technologies, Danvers, MA), mouse anti-NTPase (gift from JF Dubremetz, University of Montpellier, France) and rabbit anti-Rab11A (Cell Signaling Technologies, cat #2413S). This anti-Rab11A antibody was selected to specifically recognize human Rab11A (excluding human Rab11B and *Toxoplasma* Rab11) based on unique sequence at the C-terminus from amino acid 181 to 216 in human Rab11A. Secondary antibodies from Thermo Fisher Scientific included anti-rat conjugated to Alexa Fluor 350, Alexa Fluor 488 or Alexa Fluor 594, and anti-rabbit conjugated to Alexa Fluor 594 or Alexa Fluor 647.

### Cell and parasite cultivation

Vero cells, HeLa cells, and human foreskin fibroblasts (HFF) were obtained from American Type Culture Collection (Manassas, VA). Cells were maintained in αMEM with 10% (v/v) FBS, 100 U/mL penicillin/streptomycin (Quality Biological Inc, Gaithersburg, MD) and 2 mM L-glutamine (complete αMEM culture medium) at 37°C and 5% CO_2_. The Vero cells stably expressing GFP-Rab11A (Romano et al., 23017) was cultivated in complete αMEM culture medium containing 800 µg/mL G418. *Toxoplasma* tachyzoites (type 1 RH), WT and transgenic strains, were serially passaged in 6-well plates on confluent monolayers of HFF by transferring the supernatant containing egressed tachyzoites from one well to another (Roos *et al*., 1995). The RFP expressing RH strain was provided by F. Dzierszinski (McGill University, Montreal, Canada; Dzierszinski *et al*., 2004). The RH strain with *gra2* and *gra6* knocked out (Δgra2Δgra6) was provided by MF Cesbron-Delauw (Université Grenoble Alpes, Grenoble, France; Mercier *et al*., 2002).

### Toxoplasma infection of mammalian cells

Transfected cells were infected 4 h following transfection and untransfected cells were infected approximately 24-48 hours after plating at 60% confluency. For infection, media was removed from the cells on coverslips in wells from a 24-well plate, and freshly egressed parasites were added for 30 min. The coverslips were then washed with prewarmed PBS to remove extracellular (non-invading) parasites allowing for synchronization of infection. The infections proceeded for 24 h to yield 4 to 8 parasites per PV.

### Plasmids

The plasmids GFP-FIP1C, GFP-FIP1C^I621E^, GFP-FIP2, GFP-FIP2^I480E^, GFP-FIP3, GFP-FIP3^I738E^, GFP-FIP4, GFP-FIP4^YMD617-619AAA^, GFP-FIP5 and GFP-FIP5^I630E^ were generously provided by M. McCaffrey (University College Cork, Cork, Ireland). Their sequences were verified using the standard CMV-For primer and our primer oJR81 5’ GGGAGGTGTGGGAGGTTTT 3’. The Arf6-mCherry plasmid pcDNA3/hArf6(WT)-mCherry was a gift from Kazuhisa Nakayama (Addgene plasmid # 79422; http://n2t.net/addgene:79422; RRID:Addgene_79422; Makyio *et al*., 2012) and the sequence was confirmed using standard CMV-For and BGH-Rev primers. The plasmids containing the Arf6-binding domain(ABD) deletion mutant (GFP-FIP3-ABD) and the ABD and Rab11-binding domain (RBD) mutant (GFP-FIP3-I738E-ABD) were engineered from the McCaffrey plasmids GFP-FIP3 and GFP-FIP3^I738E^ using the Q5 Mutagenesis Kit (New England Biolabs, Ipswich, MA) following manufacturer instructions. To generate FIP3-ABD mutant, amino acids 687-712 were deleted using primers ABDdlt_F 5’ AGCTCCGTCTCCCGAGAT 3’ and ABDdlt_R 5’ CCCGTTCAGCTCCTCGTT 3’. The same pair of primers was used for both pEGFP-c1-FIP3 and pEGFP-c1-FIP3 2-756 I738E using an annealing temperature of 68°C with 3% DMSO added to the PCR reaction. Transformed bacteria were grown on LB-agar plates containing kanamycin overnight and then cultured in 3ml LB Broth with kanamycin for 22 h at 37°C at 220 rpm. After transformation and overnight culture, the Qiaprep Spin Miniprep Kit (Qiagen, Germantown, MD) was used to isolate the plasmid, and Sanger sequencing using primer oJR81 was used to confirm the sequence of the FIP3-ABD and FIP3-RBD-ABD mutant plasmids.

### Mammalian cell transfection

Vero cells were transfected with 2 µg plasmid DNA in the Amaxa Nucleofector solution R (Lonza Bioscience, Rockville, MD), following the manufacturer’s instructions with program V-01. Transfected Vero cells were then plated on coverslips in 24-well plates and allowed to recover for 24 h before infection or fixation. HeLa cells were plated on coverslips in 24-well plates at the confluency of 45-50% 24-48 h before transfection using jetPrime (Polyplus-transfection, New York, NY) for 4 h following manufacturer’s instructions. Recommended DNA and reagents for HeLa cells were adjusted to 0.2 µg plasmid DNA, 0.4 µL jetPrime reagent and 50 µL jetPrime buffer per well to minimize cell toxicity.

### Transcriptome sequencing

*Toxoplasma*-infected HFF for 24 h and uninfected HFF were PBS-washed, detached with trypsin, collected by centrifugation, and resuspended in PBS buffer. Total RNA from infected and uninfected HFF from 3 independent preparations was extracted using a Qiagen RNeasy kit. The total RNA was shipped to Novogene Corporation (Sacramento, CA) to be processed and analyzed. First, total RNA was converted to sequencing read libraries using the NEB Next Ultra kit (Illumina). The libraries were subjected to paired-end sequencing using the Novaseq 6000 with a read length of 150 bp. Each sample was sequenced to a depth of at least 20 million reads. The sequencing reads per sample were trimmed and mapped to the human genome (*Homo sapiens* GRCh38/hg38). Differential gene expression analysis was performed using DESeq R (1.18.0) and p-values were adjusted using the Benjamini and Hochberg’s approach for controlling false discovery rate (p < 0.05 considered as differentially expressed).

### Immunofluorescence Assay (IFA)

Cells were fixed in solution of 4% formaldehyde (Electron Microscopy Sciences, Hatfield, PA) and 0.02% glutaraldehyde (EMS) for 15 min at room temperature and permeabilized with 0.3% Triton X-100 for 5 minutes at room temperature. Cells were blocked with 3% Bovine Serum Albumin (BSA) for 1 h at room temperature before addition of the primary antibody diluted in 3% BSA for 1 h at room temperature or overnight at 4°C. Secondary antibodies diluted in 3% FBS were added to the coverslip for 1 h incubation at room temperature. To stain nuclei, coverslips were incubated with DAPI at a 1 µg/ml in water for 5 min at room temperature. Coverslips were then mounted with the Prolong Gold or Diamond antifade mounting solution.

### Fixed cell imaging and analysis

Cells were imaged on a Zeiss Axioimager M2 fluorescent microscope with a z motor for acquisition of slices along the *z*-axis. An oil-immersion Zeiss plan Apo 100x/NA 1.4 objective and Hamamatsu ORCA-R2 camera were used to acquire images with optical z-slices of 0.2 µm. Images were acquired, registry corrected, deconvolved, and brightness/contrast adjusted using the Volocity imaging software version 6.3 or 6.3.1 (Perkin Elmer, Waltham, MA). Fluorescent foci in the PV were identified and counted manually after identifying the boundaries of the PV based on the change in fluorescent patterns of the fluorescently tagged proteins. Statistical differences were determined using unpaired, two-tailed T-tests with the Welch’s correction for unequal standard deviations with Prism (Graphpad, San Diego, CA). PV were ranked as positive based on one or more distinct fluorescent foci detected within the bounds of the PV or negative if no foci could be observed inside the PV.

### ImmunoElectron microscopy

For immunogold staining of GFP-FIP3 and GFP-FIP3-RBD, *Toxoplasma*-infected cells were fixed in 4% paraformaldehyde (Electron Microscopy Sciences) in 0.25 M HEPES (ph7.4) for 1 h at room temperature, and then in 8% paraformaldehyde in the same buffer overnight at 4°C. They were infiltrated, frozen and sectioned as described (Romano *et al*., 2017). The sections were immunolabeled with antibodies against GFP (Clontech Laboratories Inc, Mountain View, CA) at 1:25 diluted in PBS/1% fish skin gelatin, and then with secondary IgG antibodies coupled to 10 nm protein A-gold particles before examination with a Philips CM120 EM (Eindhoven, the Netherlands) under 80 kV.

## Acknowledgements

We thank the members of the Coppens’ laboratory for helpful discussion during the course of this work and especially Karen Ehrenman for her help for the cloning of the ABD mutant. We are also grateful to the generous providers of antibodies, plasmids and parasite strains used in this study. We thank the excellent technical staff and Kim Zichichi of the Electron Microscopy Core Facility at Yale School of Medicine. This study was supported by the grant from the NIH AI060767 to I.C. and a pre-doctoral fellowship from the American Heart Association to E.J.H.

## Main abbreviations

ABD: Arf6-binding domain

ERC: endocytic recycling compartment

FIP: Rab11-Family Interacting protein

IVN: intravacuolar network

PV: parasitophorous vacuole

RBD: Rab11-binding domain

RE: recycling endosomes

SE: sorting endosomes

TfR: transferrin receptor

## Supplemental information

**Figure S1.**
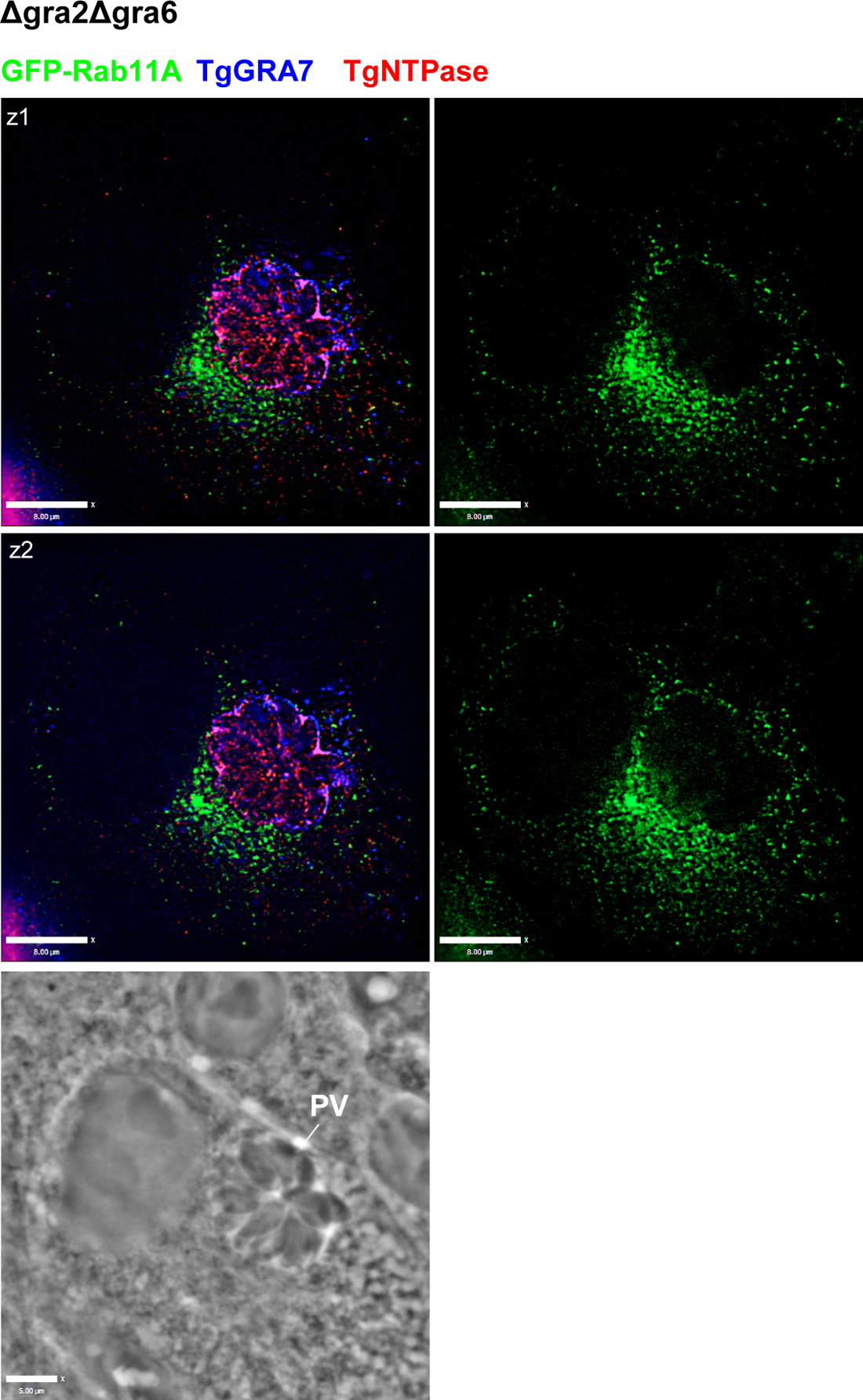
Distribution of Rab11A vesicles cells in infected cells with Δgra2Δgra6 mutants Fluorescence microscopy of Rab11A-expressing VERO cells infected with Δgra2Δgra6 mutants for 24h, before immunostaining with anti-TgGRA7 (PVM/IVN) and TgNTPase (PV lumen) antibodies.

**Figure S2.**
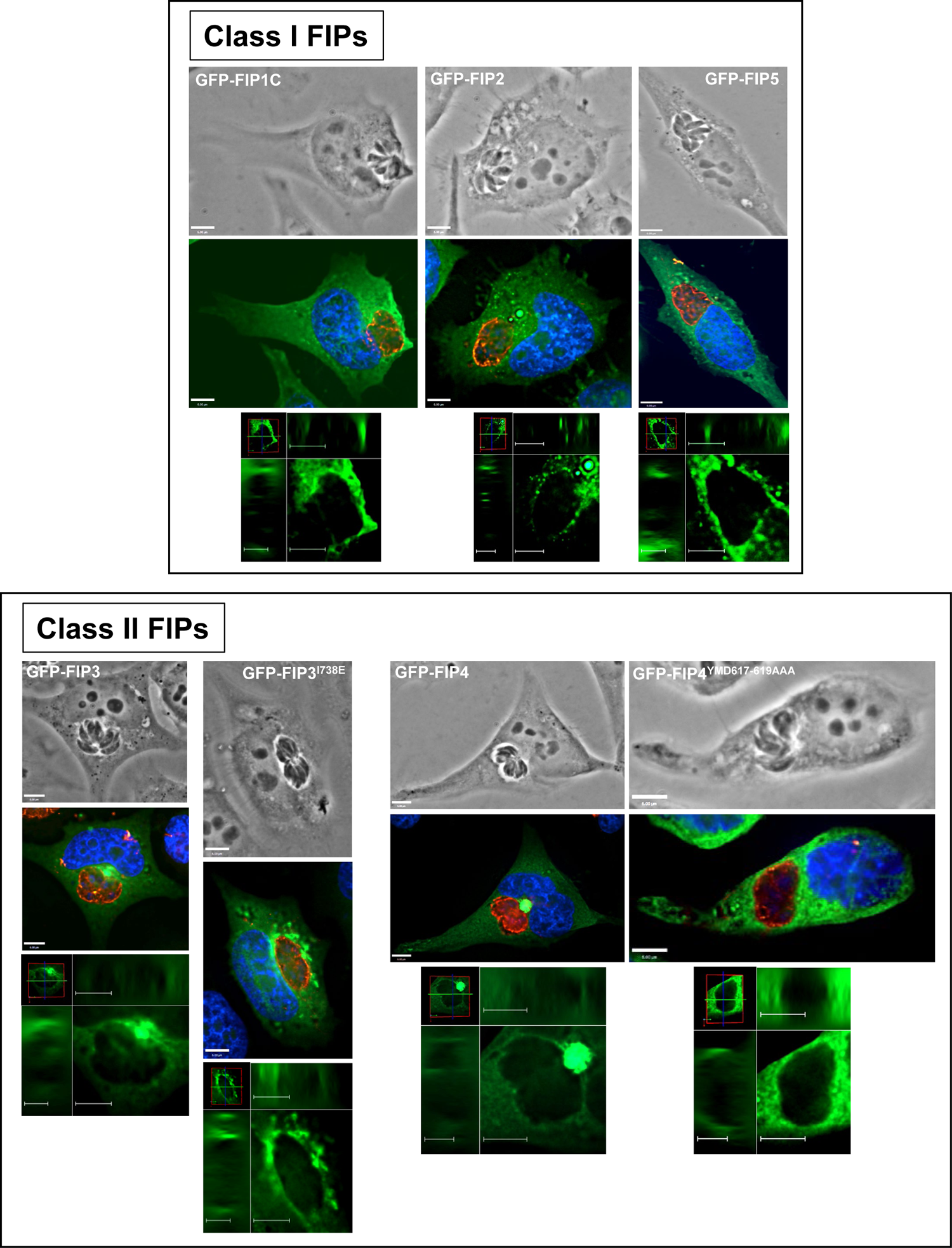
Distribution of FIP vesicles cells in infected cells with Δgra2Δgra6 mutants Fluorescence microscopy of HeLa cells transfected with GFP-FIP1C, GFP-FIP2, GFP-FIP5, GFP-FIP3, GFP-FIP3^I738E^, GFP-FIP4 or GFP-FIP4^YMD617-619AAA^, infected with *Toxoplasma* for 24h before immunostaining with anti-TgGRA7 antibody. DAPI in blue. For all images, individual z-slices are shown.

**Figure S3.**
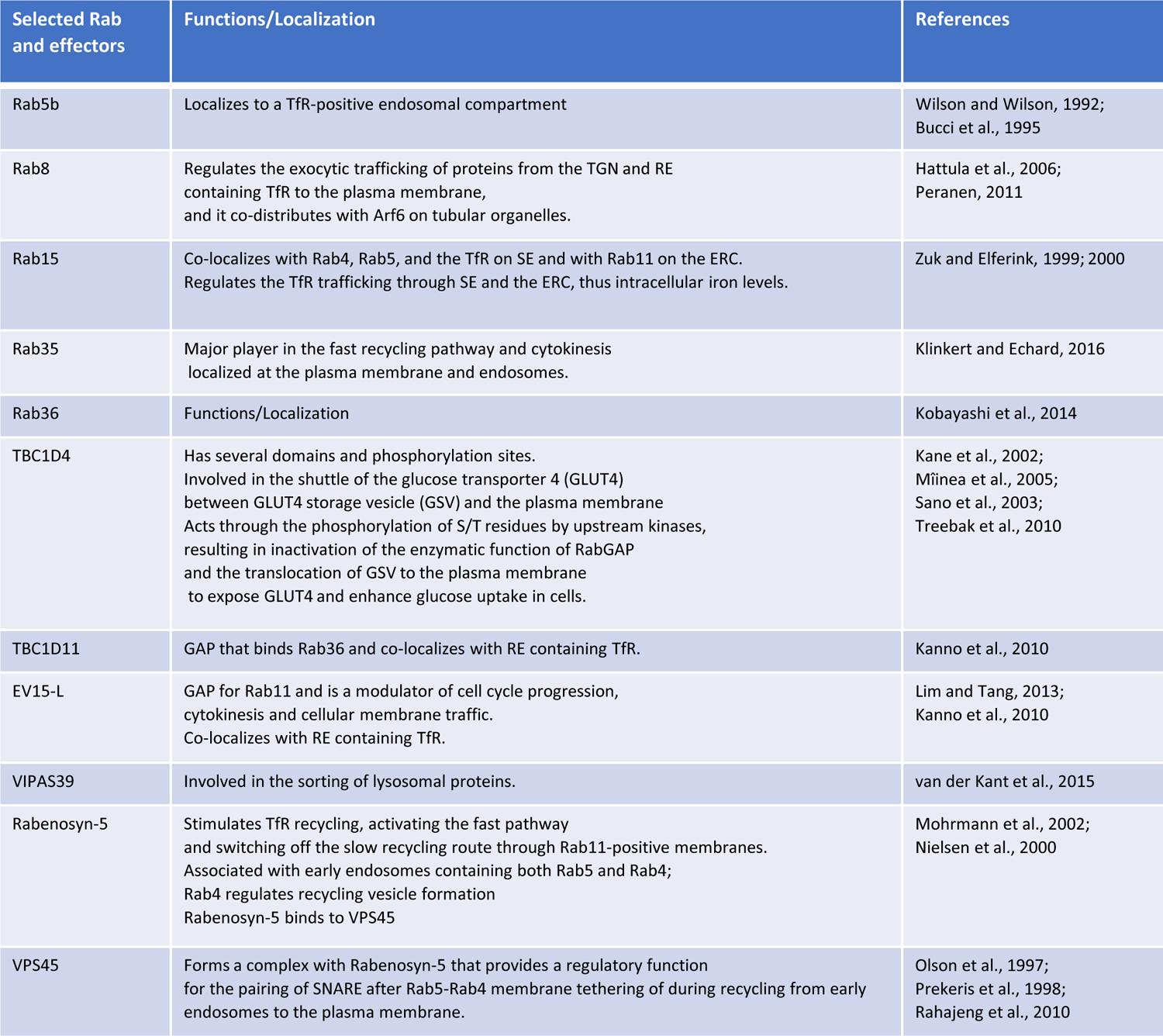
Description of selected Rabs and their effectors genes with differential expression upon *Toxoplasma* infection

**Table SI.**
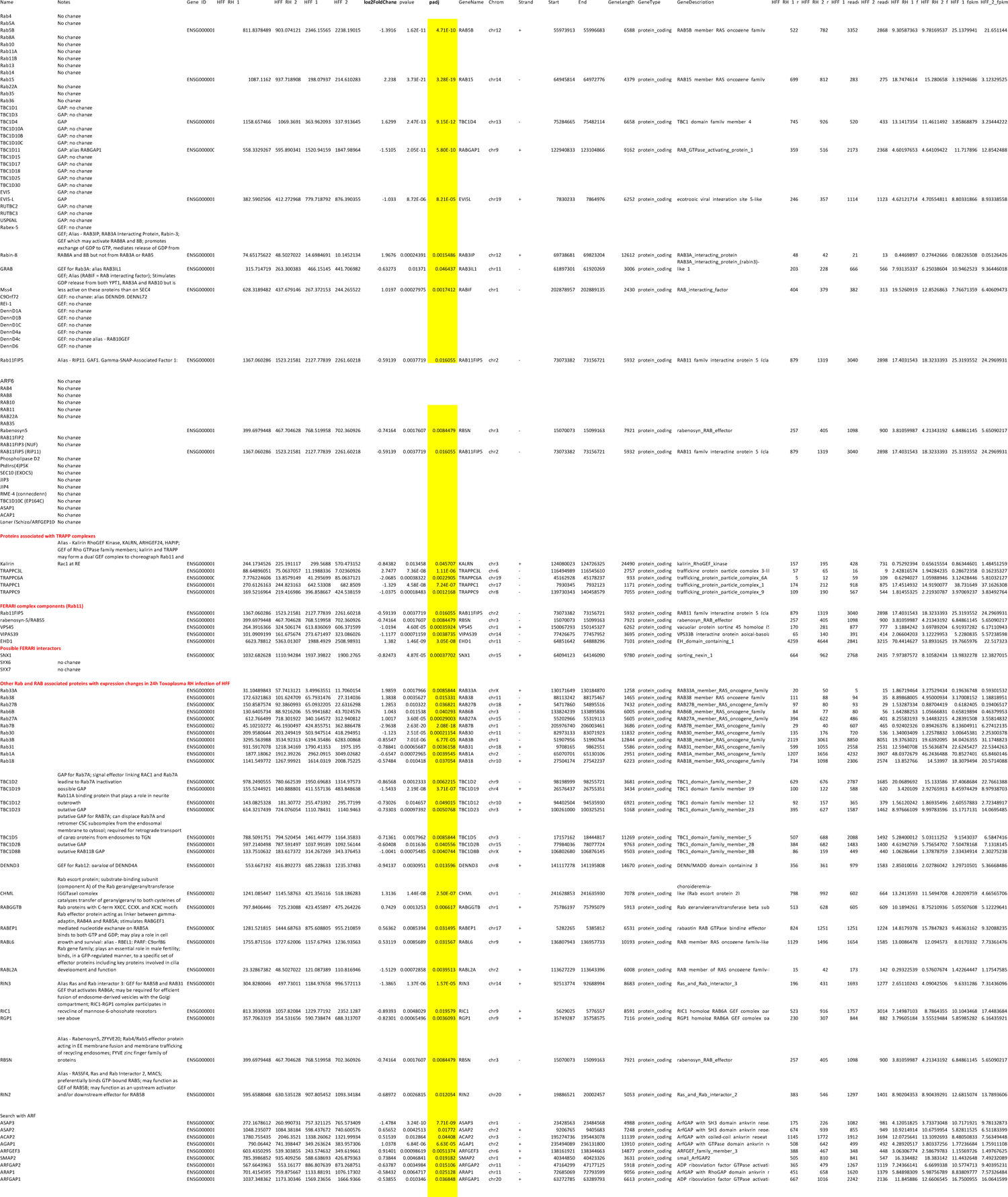
List of genes with differential expression upon *Toxoplasma* infection

